# Probabilistic modeling of personalized drug combinations from integrated chemical screen and molecular data in sarcoma

**DOI:** 10.1101/396358

**Authors:** Noah E. Berlow, Rikhi Rikhi, Mathew N. Geltzeiler, Jinu Abraham, Matthew N. Svalina, Lara E. Davis, Erin Wise, Maria Mancini, Jonathan Noujaim, Atiya Mansoor, Michael J. Quist, Kevin L. Matlock, Martin W. Goros, Brian S. Hernandez, Yee C. Doung, Khin Thway, Tomohide Tsukahara, Jun Nishio, Elaine C. Huang Huang, Susan Airhart, Carol J. Bult, Regina Gandour-Edwards, Robert G. Maki, Robin L. Jones, Joel E. Michalek, Milan Milovancev, Souparno Ghosh, Ranadip Pal, Charles Keller

## Abstract

Cancer patients with advanced disease exhaust available clinical regimens and lack actionable genomic medicine results, leaving a large patient population without effective treatments options when their disease inevitably progresses. To address the unmet clinical need for evidence-based therapy assignment when standard clinical approaches have failed, we have developed a probabilistic computational modeling approach which integrates sequencing data with functional assay data to develop patient-specific combination cancer treatments. This computational modeling approach addresses three major challenges in personalized cancer therapy, which we validate across multiple species via computationally-designed personalized synergistic drug combination predictions, identification of unifying therapeutic targets to overcome intra-tumor heterogeneity, and mitigation of cancer cell resistance and rewiring mechanisms. These proof-of-concept studies support the use of an integrative functional approach to personalized combination therapy prediction for the population of high-risk cancer patients lacking viable clinical options and without actionable DNA sequencing-based therapy.

## INTRODUCTION

Despite decades of advancements in cancer treatment, over 600,000 patients with solid tumors die annually in North America^1^, including approximately 5,000 sarcoma-related deaths. The population of high-risk, late-stage, recurrent, rare or refractory cancer patients who have exhausted standard clinical pathways and lack further treatment options represents a major unmet clinical need. Currently, DNA sequencing of tumors for druggable mutations leaves approximately 60% of patients without an actionable result^2,3^. Additionally, in many cases, single drug therapy fails to provide sustainable disease control^4^. A critical missing element in personalized cancer therapy design is the lack of effective methodologies and model-based prediction, design, and prioritization of patient-specific drug *combinations*, especially in the presence of limited tumor tissue material.

Numerous approaches to computational modeling of drug sensitivity and therapy assignment exist, in part to address ambiguity in DNA sequencing results^2,5^. These approaches are primarily based on gene expression^6^, or a combination of genomic and epigenomic data^7^. For instance, 1) integrative genomic models using Elastic Net regression techniques have been developed from large datasets such as the Cancer Cell Line Encyclopedia (CCLE)^8^ database; 2) integrative models using Random Forests with Stacking^9,10^ to integrate multiple genetic data sets for sensitivity prediction; and 3) a team science based sensitivity prediction challenge produced independent models integrating multiple data types for sensitivity prediction^11^; despite 44 individual models and a “wisdom of crowds” approach merging the top-ranked predictive models together, none of the approaches surpassed 70% predictive accuracy^11^ falling short of a reasonable accuracy threshold for clinical utility. Some recent work has focused on the use of functional data for therapy selection, such as 1) the use of microfluidics to test multiple drugs efficiently on primary patient samples^12^, 2) the use of shRNA libraries to predict drug combinations for heterogenous tumor populations^13^, and 3) a re-analysis of the CCLE database used machine learning models integrating functional response data to improve sensitivity prediction accuracy over molecular data-based Elastic Net models^14^. Integration of functional data may improve overall predictive accuracy over solely molecular data-based predictive models, especially for individual patient samples, emphasizing the need for improved drug sensitivity prediction to enable patient-specific therapy design.

To address the need for accurate prediction of drug sensitivity and design of multi-drug combinations, we previously developed a functional drug sensitivity-based modeling approach termed Probabilistic Target Inhibition Maps (PTIMs)^14–17^. The base PTIM methodology integrates quantified drug-target inhibition information (EC_50_ values) and log-scaled experimental drug sensitivities (IC_50_ values) to identify mechanistic target combinations explaining drug sensitivity data. PTIM modeling improved predictive accuracy over Elastic Net models from the CCLE dataset^14^, and has guided *in silico* validation experiments from primary canine osteosarcoma cell models^14,16–18^ and *in vitro* validation experiments^19^ on diffuse intrinsic pontine glioma (DIPG) cell models.

The key PTIM modeling assumption is that *in vitro* drug sensitivity in cancer cells is driven by a small subset of key gene targets uniquely determined by the patient’s biology, and that patient-specific drug sensitivity is most accurately predicted by multivariate modeling of autologous drug sensitivity data. The PTIM pipeline requires drug screening data from multiple (60+) monotherapy agents with quantified drug-target EC_50_ values (Fig. 1, Testing Step). PTIM modeling specifically takes advantage of the promiscuity of targeted compounds by incorporating main-target and off-target EC_50_ values during modeling. Integration of additional patient-specific molecular data (*e.g.*, exome-seq, RNA-seq, phosphoproteomics, siRNA-mediated gene knockdown, Fig. 1, Testing Step) identifies targets of interest to further refine target selection during model creation. Drug sensitivity data and secondary molecular data are provided as inputs to the PTIM computational framework^14–19^, which provides as output a mathematical model quantifying expected sensitivity of multi-target inhibition of the patient’s cancer cells. Multi-target sensitivity mechanisms are represented graphically as “tumor cell survival circuits” (Fig. 1, Modeling Step) where target combinations are denoted as “blocks” (*e.g.* Fig. 1, Modeling Step inhibitor symbols A, B, C + D). The value in the center of each PTIM block represents expected scaled sensitivity following inhibition of associated block targets. The resulting PTIM model enables combination therapy assignment via matching of targets in high-sensitivity PTIM blocks to drugs in clinical investigation or clinical use. A single block denotes monotherapy (e.g. A, B) or combination therapy (synergistic targets, e.g. C + D), while multiple blocks represent independent treatments which can be leveraged to abrogate cancer cell resistance. If PTIM models from spatially-distinct tumor sites are available, consensus therapy can be selected from distinct models to mitigate potential intra-tumor heterogeneity. When available, additional patient tumor tissue can be used to validate PTIM-predicted combination therapy *in vitro* or *in vivo* (Fig. 1, Validation Step). PTIM modeling is the foundation of our personalized therapy pipeline built to address the unmet clinical needs of the 600,000 patients dying from cancer every year^1^.

**Figure 1.**
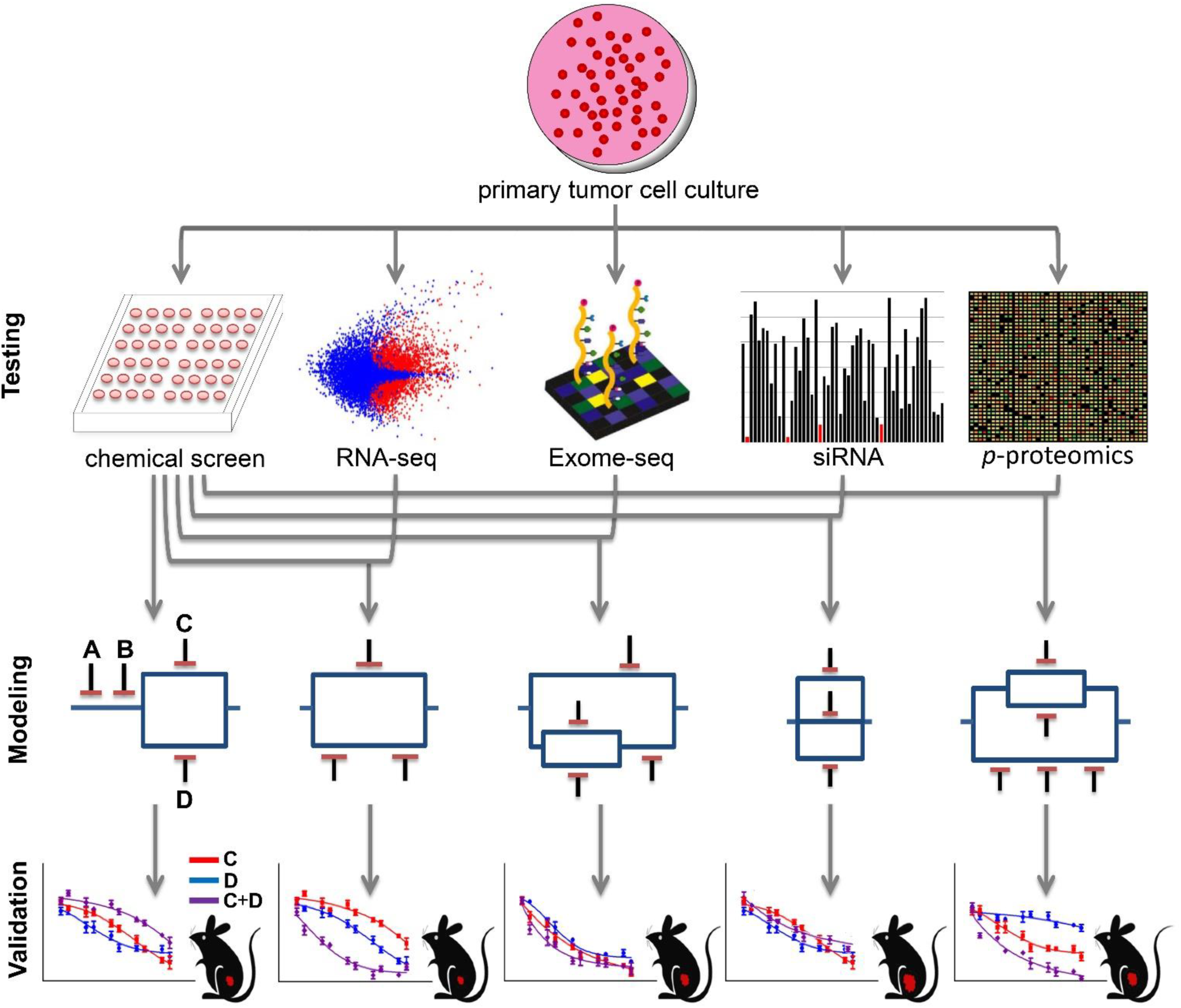
Schematic representation of experimental and computational approach to personalized combination targeted therapy predictions. Following tumor extraction and culture establishment, biological data is generated (*e.g.*, chemical screening, transcriptome sequencing, exome sequencing, siRNA interference screening and phosphoproteomic analysis) and used as input for PTIM modeling. To briefly explain the graphical model representation, targets A and B denote two independent single points of failure. Targets C and D denote parallel targets, which independently are not predicted to be effective, but together will be synergistic and lead to significant cell growth inhibition. Targets A, B, and the C-D parallel block are in series and may target independent pathways. Series blocks, when inhibited together, may abrogate cancer resistance mechanisms by knockdown of independent pathways. Model sensitivity scores then guide *in vitro* validation and *in vivo* validation experiments.

Herein, we present proof-of-concept validation experiments of the integrative PTIM pipeline (Fig. 1) using soft tissue sarcoma as a paradigm. Each validation experiment applies PTIM combination therapy design to address one of three critical unmet needs in cancer treatment: 1) selection of functional evidence-based synergistic drugcombinations, validated inmurine alveolar rhabdomyosarcoma (aRMS);2) consensus modeling of multi-site drug sensitivity data to overcome intra-tumor heterogeneity, validated in epithelioid sarcoma (EPS); and 3) resistance abrogation by targeting of parallel biological pathways, validated in undifferentiated pleomorphic sarcoma (UPS).

## RESULTS

### Proof-of-concept of synergy prediction by PTIM modeling

#### Chemical screening, biological interrogation, and PTIM modeling of a GEMM-origin aRMS

For our 2-drug synergy proof-of-concept study, we used a low passage primary tumor cell culture of a GEMM-origin aRMS tumor designated U23674^20^ as a pilot study of the PTIM personalized therapy pipeline. From our previous work^21,22^ we reasoned that kinases would be fundamental to the biology of aRMS, thus we interrogated U23674 drug sensitivity via three kinase inhibitor compound libraries: the GlaxoSmithKline (GSK) Open Science Orphan Kinome Library (GSK screen), the Roche Orphan Kinome Screen Library (Roche screen), and a custom Pediatric Preclinical Testing Initiative Drug Screen Version 2.1 (PPTI screen).

The GSK screen^23^ consists of 305 compounds with experimentally quantified drug-target interaction EC_50_ values. Of the 305 screened compounds, 40 (13%) caused at least 50% cell growth inhibition at or below maximum tested *in vitro* dosage in in U23674, hereafter defined as a compound “hit” (Supplemental Fig. S1 and Supplemental Tables S1 and S2). The Roche screen consists of 223 novel kinase inhibitor compounds, most with quantified drug-target interactions; 21 of 223 compounds (9.4%) were hits on U23674 (Supplemental Fig. S2 and Supplemental Tables S3, S4 and S5). The PPTI screen consists of 60 preclinical‐ or clinical-stage targeted agents; 28 of 60 compounds (46.7%) were hits on U23674 (Supplemental Fig. S3 and Supplemental Tables S6 and S7).

Additionally, primary tissue was sequenced (tumor whole exome sequencing, matched normal whole exome sequencing, and whole transcriptome sequencing, Supplemental Tables S8 and S9). Exome sequencing of U23674 did not identify any druggable targets both mutated and amplified (Supplemental Fig. S4 and Supplemental Tables S8 and S9); six genes possessed activating mutations (*Fat4, Gm156, Mtmr14, Pcdhb8, Trpm7, Tttn, Zfp58*) and one gene possessed a high-impact frameshift indel (*Ppp2r5a*); none of these seven gene targets are druggable. No gene with a mutation or indel is druggable. Four druggable gene targets show evidence of copy number gain (*Gsk3a, Epha7, Psmb8, Tlk2*). *Gsk3a*, *Psmb8*, and *Tlk2* all show neutral expression or underexpression by RNA-seq. *Gsk3a* inhibitors were effective in 12 of 72 inhibitors (16.667%) across three screens, suggesting Gsk3a is not critical for cancer cell survival. *Psmb8* inhibition showed *in vitro* efficacy in nearly all tested cell cultures across multiple tumor types (unpublished internal data) and, along with lack of overexpression, was thus treated as an *in vitro* screening artifact; furthermore, clinical response of solid tumors to proteasome inhibitors has been limited^24^. *Tlk2* has no published inhibitor compounds. While overexpressed, the *Epha7* inhibitor on the PPTI drug screen was ineffective against U23674. Therapy assignment via exome sequencing would thus have limited clinical utility for U23674.

#### *Probabilistic Target Inhibition Map (PTIM) modeling identifies 2-drug combinations with synergy* in vitro

The high average level of target coverage (24 compounds/target), the inclusion of both typical and atypical kinase target combinations, and the thorough characterization of drug-target interactions made the GSK screen the most complete dataset available and was thus selected to guide *in vitro* and *in vivo* validation experiments. Baseline (chemical screen data only), RNA-seq informed-, exome-seq informed-, siRNA interference informed-, and phosphoproteomics informed‐ PTIM models were generated from the GSK screen data (Fig. 2A-C, Supplemental Fig. S5, Supplemental Tables S10 - S12). PTIM-identified targets were consistent with known targets of interest in aRMS^25,26^ and identified gene targets involved in established protein-protein interactions^27^ (Supplemental Fig. S6). As multi-drug combinations impart toxicity concerns and dosing limitations, we focus on PTIM blocks (combinations of two or more targets) treatable by at most two drugs. Baseline and genomics-informed PTIM models were also generated for the PPTI and Roche screens (Supplemental Fig. S7), however no validation experiments based on PPTI or Roche PTIM models were performed due to focus on the GSK screen results.

**Figure 2.**
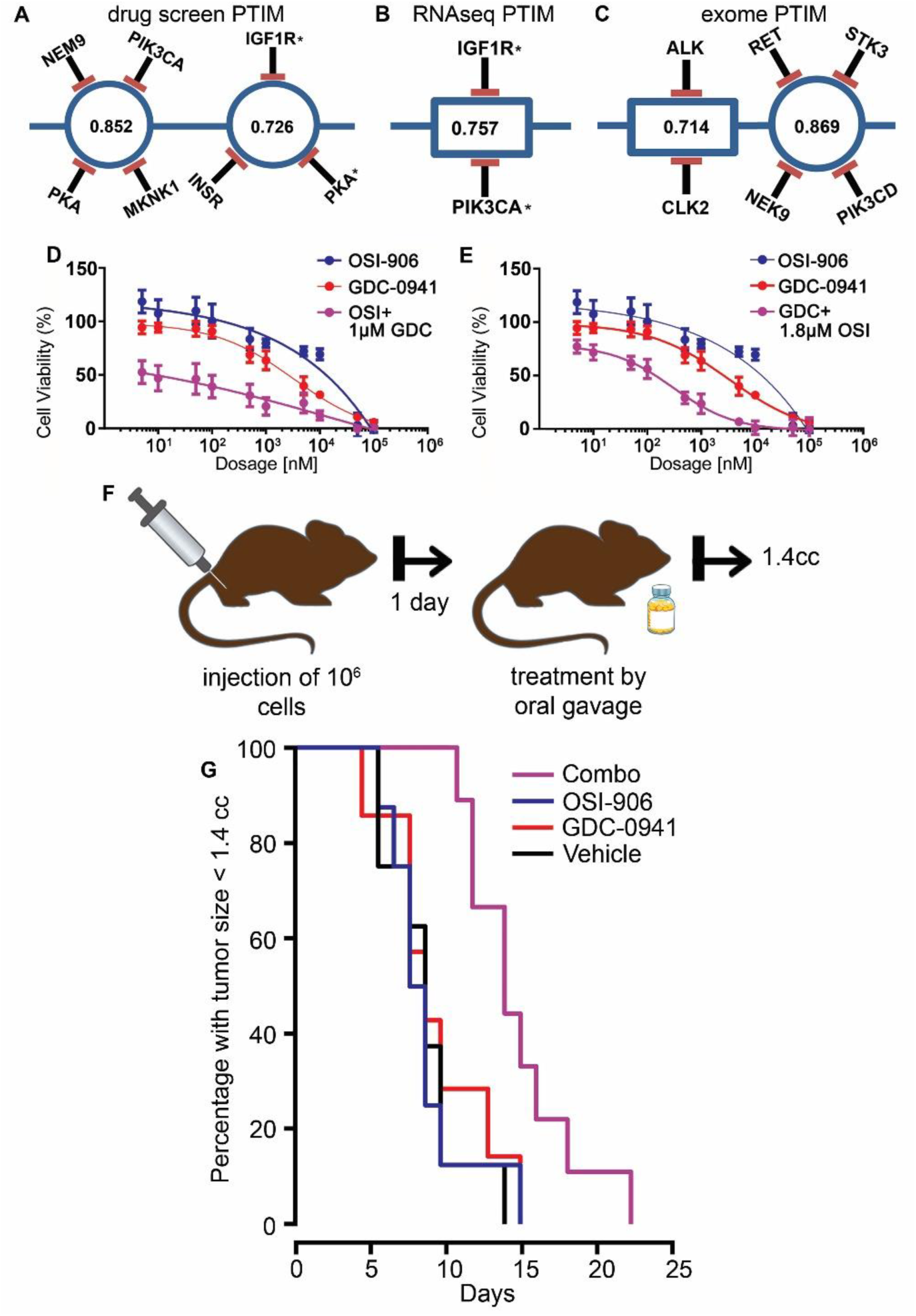
Probabilistic Target Inhibition Maps (PTIMs) and experimental *in vitro* and *in vivo* results for U23674 alveolar rhabdomyosarcoma (aRMS) drug combinations. Targets with adjacent asterisks indicate targets selected for *in vitro* validation. Values in the center of PTIM blocks represent expected scaled sensitivity following inhibition of associated block targets. (**A**) Abbreviated chemical screen-informed PTIM. (**B**) Abbreviated RNA-seq + Chemical screen informed PTIM. (**C**) Abbreviated exomeseq + Chemical screen informed PTIM. The values within the target blocks indicate scaled drug sensitivity for the given target combination^16^ when the targets are inhibited via one or more chemical compounds. More information can be found in prior publications^16,18^. In (**D**–**E**), results are based on n = 3 independent experiments with n = 4 technical replicates. (**D**) Dose response curve for OSI-906 varied dosage + GDC-0941 fixed dosage. The response for GDC-0941 at varied dosages is included. (**E**) Dose response curve for GDC-0941 varied dosage + OSI-906 fixed dosage. The response for OSI-906 at varied dosages is included. (**F**) Schematic representation of *in vivo* experiment design. (**G**) Kaplan-Meier survival curves for *in vivo* orthotropic mouse experiment. Mice were treated with vehicle (n = 8 mice, black line), 50 mg/kg OSI-906 (n = 8 mice, blue line), 150 mg/kg GDC-0941 (n = 7 mice, red line), or combination 50 mg/kg OSI-906 + 150 mg/kg GDC-0941 (n = 8 mice, purple line).

We selected one combination for *in vitro* synergy validation: RNA-seq-informed PTIM-derived target combination *Igf1r* & *Pik3ca* (Fig. 2B) with combination therapy OSI-906 (a Igf1r and Insr inhibitor) + GDC-0941 (a Pik3ca inhibitor selective against AKT/mTOR). Both compounds were selected based solely on selectivity of interaction with the PTIM-identified targets. We selected the RNA-seq-informed drug combination due to high block sensitivity, targetability by a two-drug combination, and our previous work showing higher correlation between transcriptome status and drug sensitivity^14^. *In vitro* validation experiments for OSI-906 + GDC-0941 (Fig. 2D-E) demonstrated synergy as determined by non-constant ratio Combination Index^28^ (CI) values (Supplemental Table S13). Low-dose combination experiments were also performed to confirm PTIM-predicted drug mechanism of action (Supplemental Fig. S8, Table S13).

#### Tumor cell rewiring following synergy-focused combination therapy

To explore tumor rewiring (activation of secondary signaling pathways to improve chance of survival) following synergy-focused intervention, we treated U23674 cell populations with low-dose monotherapy or combination therapies defined in initial *in vitro* validation experiments, and subsequently screened the populations via the Roche screen (Supplemental Figs. S9 and S10 and Supplemental Table S14). Unsurprisingly, the cell populations showed evidence of rewiring within hours of monotherapy or combination therapy intervention (Supplemental Fig. S11), emphasizing the importance of simultaneous, multi-pathway drug combinations at full therapeutic doses. While PTIM modeling currently focuses on 2-drug combinations to minimize toxicity concerns, PTIM-predicted combinations of three or more drugs are possible with sufficient evidence of safety and efficacy.

Mice were treated with vehicle (n = 8 mice, black line), 50 mg/kg OSI-906 (n = 8 mice, blue line), 150 mg/kg GDC-0941 (n = 7 mice, red line), or combination 50 mg/kg OSI-906 + 150 mg/kg GDC-0941 (n = 8 mice, purple line).

#### *Probabilistic Target Inhibition Map (PTIM) modeling predicts 2-drug combination with* in vivo *efficacy*

Having demonstrated *in vitro* synergy, we next validated OSI-906 + GDC-0941 *in vivo*. We designed a four-arm orthotopic allograft study (Fig. 2F) comparing vehicle, OSI-906 (50 mg/kg), GDC-0941 (150 mg/kg), and OSI-906 (50 mg/kg) + GDC-0941 (150 mg/kg). Kaplan-Meier survival analysis (Fig. 2G) showed improvement in mouse lifespan from combination treatment (under Bonferroni correction: Vehicle – Combo, p = 0.005, OSI-906 – Combo, p = 0.014, GDC-0941 – Combo, p = 0.079. In all cases, p < 0.05 uncorrected). Survival of mice treated with either OSI-906 or GDC-0941 alone was indistinguishable from treatment by vehicle (p > 0.5, both corrected and uncorrected). Since a PTIM block represents targets which are weak independently but synergistic together, U23674 *in vivo* data supports the hypothesis underlying our modeling approach: synergistic combination targets can be identified through computational modeling of monotherapy chemical agents.

### Proof-of-concept of heterogeneity-consensus 2-drug combinations predicted by PTIM modeling

#### Development of heterogeneous cell models of Epithelioid Sarcoma (EPS)

EPS is a soft tissue sarcoma of children and adults for which chemotherapy and radiation provides little improvement in survival^29^. Effective options beyond wide surgical excision are presently undefined^30^, making EPS a viable test case for developing targeted personalized therapies.

We have developed several new heterogeneous EPS preclinical resources: three new unpublished cell cultures, as well as (to our knowledge) the first reported patient-derived xenograft (PDX) model of EPS derived from a 22-year-old female with a large proximal (shoulder) EPS tumor (Fig. 3A). The tumor sample was obtained from surgical resection and was assigned the internal identifier PCB490. Due to the size of the acquired tumor sample and the potential for heterogeneity in solid tumors^31^, we divided the 1 cm^2^ resected tumor mass into five spatially-distinct regions (designated PCB490-1 through PCB490-5) and cultured each to develop heterogeneous cell models (Fig. 3A). PCB490 cultures were maintained at low passage to minimize biological drift from the original patient samples. To confirm EPS diagnosis, three of five tumor sites (1,2, and 5) were validated by western blot for INI1 protein, which is absent in 93% of EPS samples (Fig. 3B)^29^ as well as in published cell lines^32,33^. Multiple sites were submitted to The Jackson Laboratory for establishment of PDX models; PCB490-5 PDX developed a passageable tumor that matched the original PCB490-5 sample by both histology and INI1 immunohistochemical staining (Fig. 3C–3F).

**Figure 3.**
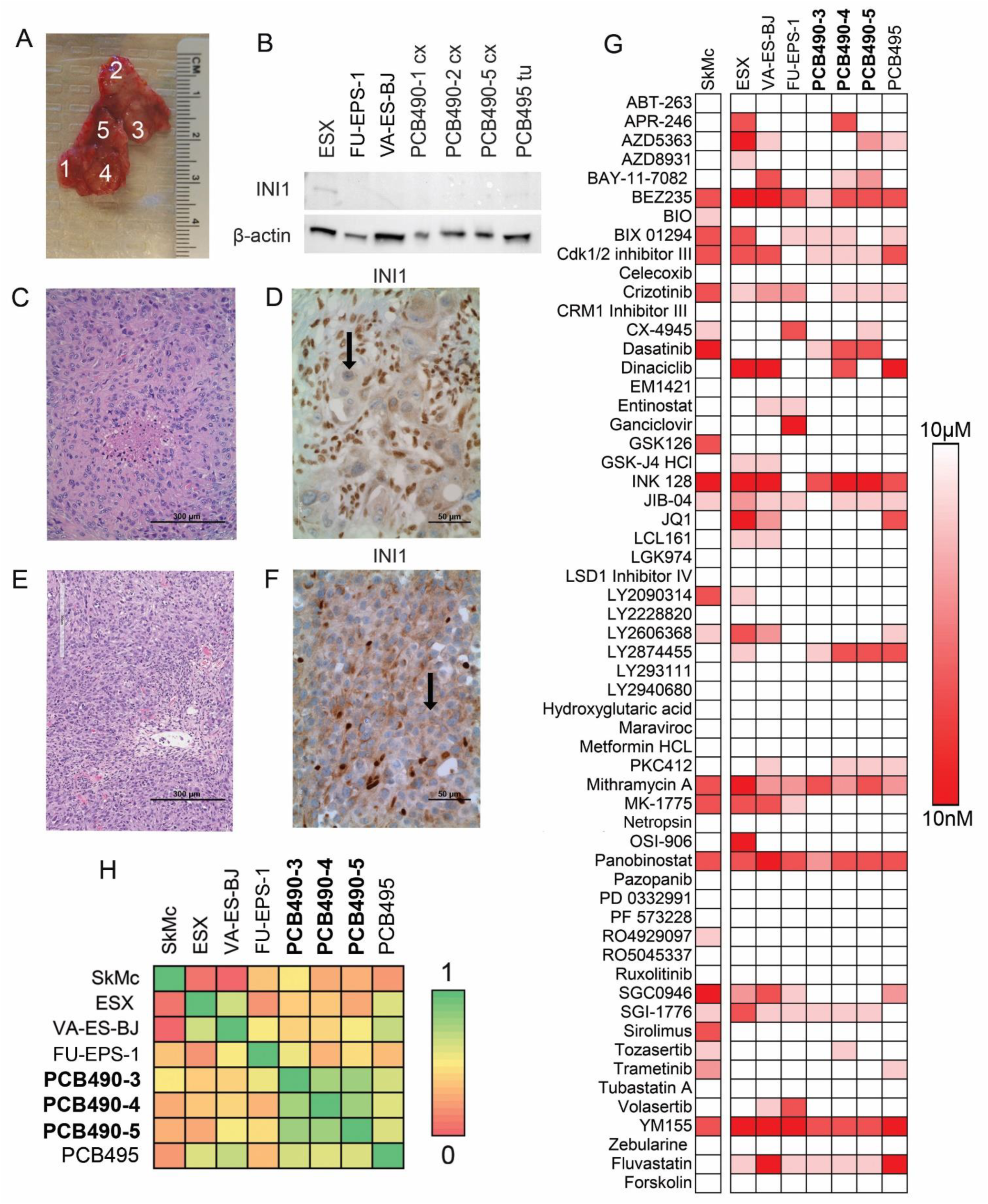
New cell cultures and patient-derived xenograft model of EPS with chemical space characterization. (**A**) PCB490 biopsy sample divided into distinct regions to create different primary tumor cell cultures for study. (**B**) Western blot demonstrating loss of INI1 in primary tumor and published EPS cell lines. (**C**) Histology of surgical biopsy. (**D**) Immunohistochemical staining for INI1 shows absence in tumor cells (black arrow) but presence in co-mingled non-cancerous cells. (**E**) Histology of PCB490 patient-derived xenograft. (**F**) INI1 absence (black arrow) in immunohistochemical staining of PCB490 patient-derived xenograft. (**G**) Drug Screen V3 results from primary EPS cell cultures, published EPS cell lines, and a normal myoblast cell line. (**H**) Heatmap of correlation coefficients of 60-agent drug screen results.

#### Drug screening, sequencing, and comparison of heterogeneous EPS cell cultures

Cell cultures PCB490-3, PCB490-4, and PCB490-5 grew to sufficient populations (minimum 3 x 10^6^ cells) at low passage (passage 2 or below) to allow for drug screening via the investigator selected 60-agent screen denoted Drug Screen V3 (Fig. 3G, Supplemental Table S15) and the previously described Roche screen (Supplemental Table S16). Drug screen endpoints were per-drug IC_50_ values.

PCB490 primary tissue was sequenced for tumor whole exome sequencing, matched normal whole exome sequencing, and whole transcriptome sequencing (Supplemental Fig. S12, Supplemental Tables S17 and S18). Sequencing identified germline and tumor amplified, expressed, high-impact druggable variations in two genes (*ABL1*, *NOTCH1*) and expressed, medium impact variations in three additional genes (*MDM4*, *PAK4*, *MAP4K5*). All five variations were identified in both tumor and normal (germline) samples. The *ABL1* variation was identified in the 1000 Genomes Project^34^. *ABL1*, *NOTCH1*, *MDM4* and *PAK4* variations were identified in dbSNP^35^. All variants are of unknown clinical significance (Supplemental Table S19)^35,36^. PCB490 drug screening results revealed no pathway-specific drug sensitivity of mutated genes (Supplemental Fig. S13) suggesting therapy assignment via exome sequencing would likely have limited clinical utility for PCB490.

To compare drug sensitivity of PCB490 with other EPS models, three cell lines (ESX, FU-EPS-1, and VA-ES-BJ), a second human-derived cell culture (PCB495), and the SkMc skeletal myoblast cell line were assayed with Drug Screen V3 (Fig. 3G, Supplemental Table S20). Drug Screen V3 responses were compared by calculating Pearson correlation coefficients (Fig. 3H) to quantify the similarity between the new EPS models and existing EPS cell models. Correlation within primary cell cultures (PCB490 sites and PCB495) was significantly higher than correlation between primary cultures and cell lines (μ = 0.7265 vs. μ = 0.5551, p < 0.01), suggesting EPS primary cultures may be biologically distinct from EPS cell lines. PCB490 drug screen response differed between sample locations millimeters away from each other, reflective of biological differences arising from spatial tumor heterogeneity. Nonetheless, correlation between chemical screen results from PCB490 cultures was significantly higher than correlation between PCB490 cultures and PCB495 cultures/EPS cell lines (μ = 0.7967 vs. μ = 0.5807, p < 0.01), suggesting that treatments for PCB490 may be better defined solely by PCB490 biological data.

#### *PTIM modeling guides heterogeneity-consensus* in vivo *drug combination*

Highly correlated yet heterogeneous PCB490 drug sensitivity data guided us towards PTIM modeling to design a heterogeneity-consensus personalized drug combination therapy for PCB490. PTIM models for 60-agent drug screens of PCB490-3 (Fig. 4A), PCB490-4 (Fig. 4B), and PCB490-5 with integrated RNA-seq data (Fig. 4C) indicated common efficacious mechanisms across the heterogeneous tumor sites: epigenetic modifiers (HDAC, EHMT), PI3K/mTOR inhibition, and VEGF (KDR) signaling inhibition. We focused on high-scoring PTIM blocks treatable by a two-drug combination, resulting in selection of BEZ235 (PI3K/mTOR inhibitor) and sunitinib (poly-kinase inhibitor, including KDR and AXL). BEZ235 + sunitinib was selected solely based on PTIM modeling data, agnostic to previous use of sunitinib in EPS^37^. To replicate potential clinical conditions for personalized combination therapy, we bypassed *in vitro* validation and directly initiated *in vivo* testing of BEZ235 + sunitinib in the PCB490 PDX model. Though the PCB490 PDX model originates from the PCB490-5 region, heterogeneity of PCB490 suggests the tumor section used to establish the PCB490 PDX can be considered a unique heterogeneous region. PDX testing of BEZ235 + sunitinib demonstrated significant slowing of tumor growth over vehicle control (92% slower tumor growth at Day 19, p=0.01) (Fig. 4D). In statistical analysis restricted to treated animals at Day 19, BEZ235 + sunitinib significantly slowed PDX tumor growth compared to both BEZ235 (p=0.01) and sunitinib (p=0.01) alone (Fig. 4D).

**Figure 4.**
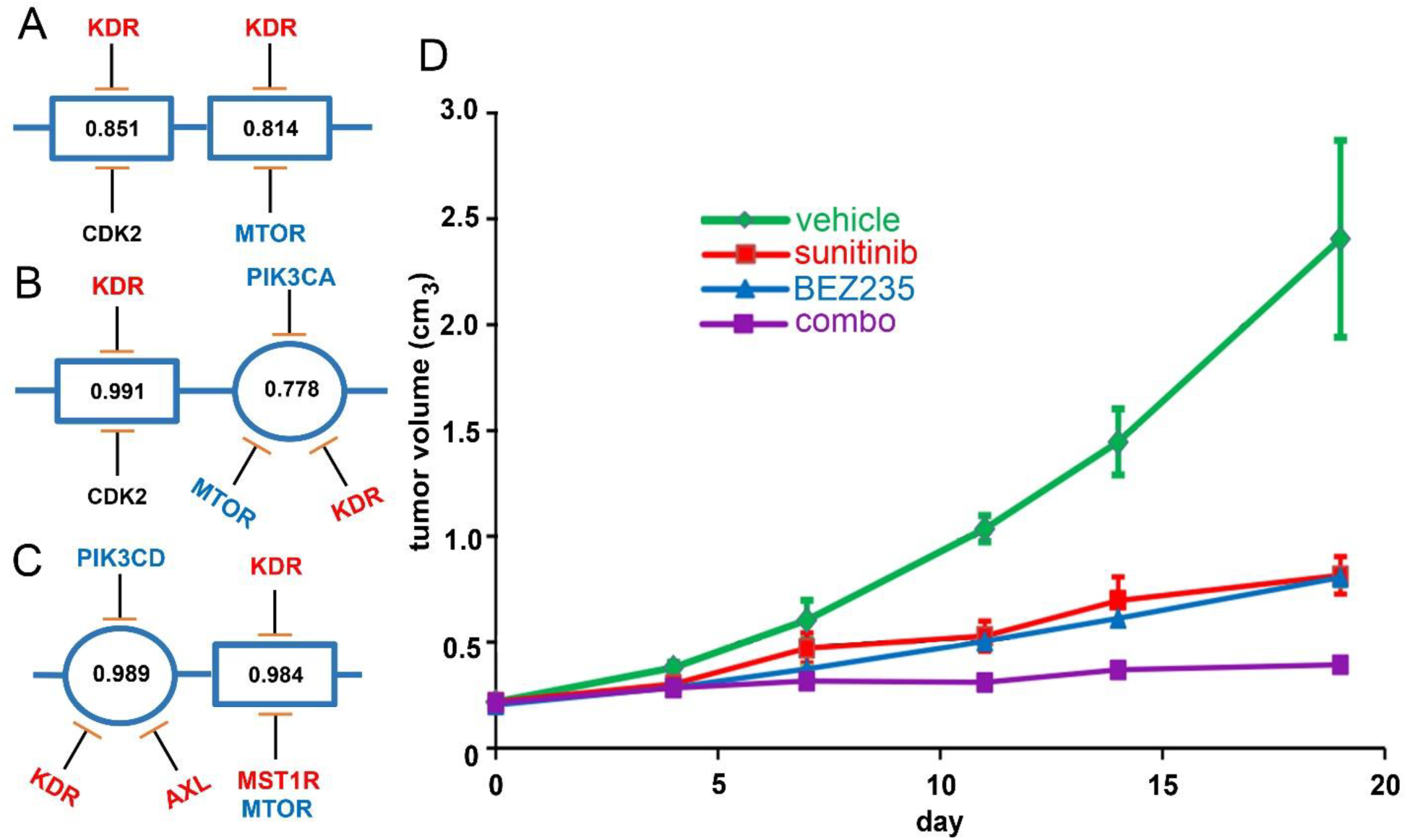
Probabilistic Target Inhibition Maps (PTIMs) of Drug Screen V3 and Roche screen results for spatially-distinct epithelioid sarcoma tumor regions. Values in the center of PTIM blocks represent expected scaled sensitivity following inhibition of associated block targets. (**A-C**) PTIMs informed by Roche Orphan Kinase Library and V3 screens. Targets of sunitinib are highlighted red, targets of BEZ235 are highlighted blue. (**A**) Abbreviated PTIM for PCB490-3. (**B**) Abbreviated PTIM for PCB490-4. (**C**) Abbreviated PTIM from PCB490-5 with integrated RNA-seq data. (**D**) Results from PCB490-5 patient-derived xenograft *in vivo* validation studies presented as group-wide tumor volumes following vehicle treatment (n = 3 mice, green line), treatment by 30.0 mg/kg sunitinib (n = 3 mice, red line), treatment by 25.0 mg/kg BEZ235 (n = 3 mice, blue line), and treatment by 25.0 mg/kg BEZ235 + 30.0 mg/kg sunitinib (n = 3 mice, purple line).

### Proof-of-concept of resistance-abrogating 2-drug combinations predicted by PTIM modeling

#### *PTIM modeling of undifferentiated pleomorphic sarcoma (UPS) samples guides cross-species resistance-abrogating drug combination* in vitro

The previously discussed U23674 rewiring experiment emphasized the need for multi-pathway targeting when developing personalized treatments. The PTIM modeling approach identifies mechanisms driving *in vitro* drug sensitivity by identifying effective target combination “blocks”; two blocks operating on different biological pathways represent two independent treatment mechanisms. We reasoned that two-block inhibition could result in resistance-abrogating combination treatments, thus we validate a drug combination built from two PTIM blocks identifying independent biological pathways. PTIM modeling of PPTI screen data from a UPS derived from a 75-year-old man (PCB197, Fig. 5A, Supplemental Table S21) and a canine-origin UPS (S1-12, Fig. 5B, Supplemental Table S21) identified species-consensus drug sensitivity mechanisms targetable by a 2-block, 2-drug combination (Fig. 5C, D): panobinostat (pan-HDAC inhibitor, HDAC7 block) and obatoclax (MCL1 inhibitor). The combination of panobinostat + obatoclax was predicted to abrogate resistance mechanisms and prevent cancer cell rewiring and regrowth; furthermore, the cross-species nature of the experiment supports the resistance-abrogation effect not being model specific.

**Figure 5.**
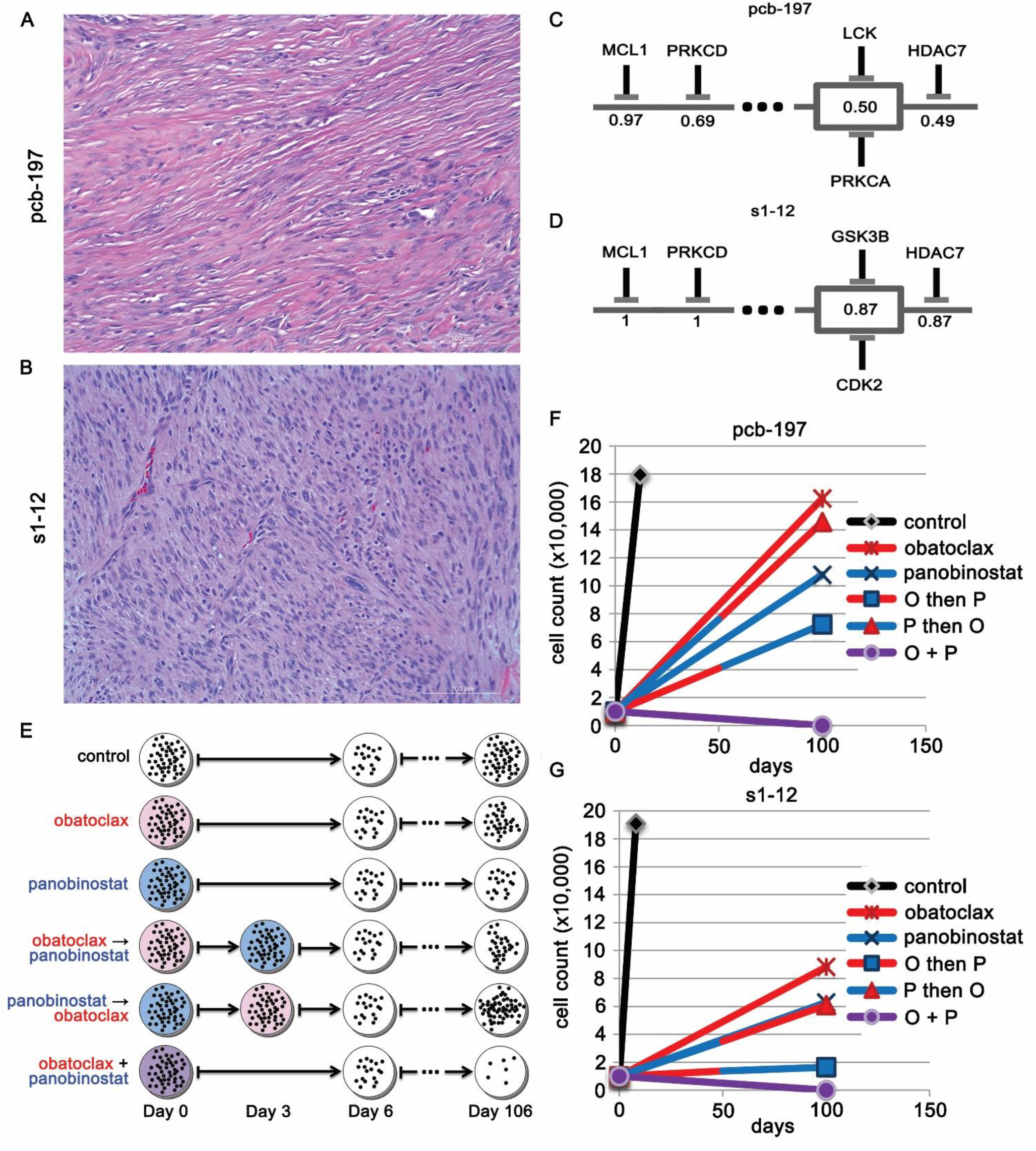
Undifferentiated pleomorphic sarcoma (UPS) Probabilistic Target Inhibition Map (PTIM)-guided resistance abrogation experiments. Values in the center of PTIM blocks represent expected scaled sensitivity following inhibition of associated block targets. (**A**) Histology of PCB197 human UPS sample (20x magnification). (**B**) Histology of S1-12 canine UPS sample (20x magnification). (**C**) Abbreviated PTIM model for the pediatric preclinical testing initiative (PPTI) screen of PCB197 human UPS sample. (**D**) Abbreviated PTIM model built from the PPTI screen of S1-12 canine UPS sample. (**E**) Schematic of experimental design for resistance abrogation experiments. (**F**) Cellular regrowth of PCB197 human UPS sample over 100 days following treatment by single and multi-agent compounds in sequence and in combination. (**G**) Cellular regrowth of S1-12 canine UPS sample over 100 days following treatment by single and multi-agent compounds in sequence and in combination. Data in (**F**–**G**) is based on n = 4 replicate experiments.

To validate *in vitro* resistance abrogation across species, we performed identical six-arm *in vitro* trials for PCB197 and S1-12. Each arm represented a different combination method for the cross-species combination: vehicle treatment, monotherapy treatment, serial monotherapy treatment (panobinostat then obatoclax, obatoclax then panobinostat), and simultaneous combination treatment (concurrent panobinostat + obatoclax) (Fig. 5E). Resistance abrogation in each arm was determined by cellular regrowth over 100 days following treatment. Rewiring and regrowth was expected for all monotherapy and serial treatment modalities. All arms except the simultaneous combination treatment arm experienced cellular regrowth, indicating the development of resistance. In both cultures, the simultaneous combination treatment arm showed no cellular regrowth over 100 days, indicating the combination potentially addressed resistance mechanisms (Fig. 5F, G).

## METHODS

### Cell model establishment

The mouse primary tumor cell culture U23674 was established from a tumor at its site of origin in a genetically engineered *Myf6Cre,Pax3:Foxo1,p53* mouse bearing alveolar rhabdomyosarcoma (aRMS) as previously described^38^. In brief, the tumor was minced and digested with collagenase (10 mg/ml) overnight at 4°C. Dissociated cells were then incubated in Dulbecco’s Modified Eagle’s Medium (DMEM) (11995-073; Thermo Fisher Scientific, Waltham, MA, USA) supplemented with 10% fetal bovine serum (FBS) (26140079; Thermo Fisher Scientific) and 1% penicillin-streptomycin (15140-122; Thermo Fisher Scientific) in 5% CO_2_ at 37°C.

The human epithelioid sarcoma (EPS) sample PCB490 was collected from a patient undergoing planned surgical resection. Tumor tissue was partitioned into 5 distinct regions, minced and digested with collagenase type IV (10 mg/ml) overnight at 4°C. The dissociated cells were then incubated in RPMI-1640 (11875-093; Thermo Fisher Scientific, Waltham, MA, USA) supplemented with 10% fetal bovine serum (FBS) and 1% penicillin-streptomycin in 5% CO_2_ at 37°C. Sections 3, 4, and 5 (PCB490-3, PCB490-4, PCB490-5) successfully grew in culture. Samples from each region were also sent to The Jackson Laboratory (JAX) for patient-derived xenograft (PDX) model establishment. Cultures were maintained at low passage to minimize biological variation from the original patient tumor. Remaining tumor pieces were snap frozen for future DNA, RNA and protein isolation.

The human EPS sample PCB495 was received through the CCuRe-FAST tumor bank program. To create the cell cultures from the PCB495 primary tumor, the tumor was minced and digested with collagenase (10 mg/ml) overnight at 4°C. The dissociated cells were then incubated in RPMI-1640 media supplemented with 10% fetal bovine serum (FBS) and 1% penicillin-streptomycin in 5% CO_2_ at 37°C.

The human undifferentiated pleomorphic sarcoma (UPS) PCB197 was received through the CCuRe-FAST tumor bank program. To create the cell cultures from the PCB197 primary tumor, the tumor was minced and digested with collagenase (10 mg/ml) overnight at 4°C. The dissociated cells were then incubated in RPMI-1640 media supplemented with 10% fetal bovine serum (FBS) and 1% penicillin-streptomycin in 5% CO_2_ at 37°C.

All human tissue samples were acquired through the Childhood Cancer Registry for Familial and Sporadic Tumors (CCuRe-FAST) tumor banking program. All patients enrolled in CCuRe-FAST provided informed consent. All aspects of the study were reviewed and approved by the Oregon Health & Science University (OHSU) Institutional Review Board (IRB). Patient data and clinical and pathologic information are maintained in a de-identified database.

The canine UPS sample S1-12 was obtained from Oregon State University’s (OSU) College of Veterinary Medicine. OSU Institutional Animal Care and Use Committee (IACUC) approval was obtained for procurement of the tissue. To establish S1-12 cell culture, tumor tissue was minced and digested with collagenase (10 mg/ml) overnight at 4°C. The dissociated cells were then incubated in RPMI-1640 media supplemented with 10% fetal bovine serum (FBS) and 1% penicillin-streptomycin in 5% CO_2_ at 37°C.

### Immunoblotting of PCB490

Tumor tissue and cells from PCB490-1,2, and 5 were lysed in radioimmunoprecipitation (RIPA) buffer containing both protease and phosphatase inhibitors (Sigma Aldrich, St. Louis, MO). Lysates were homogenized and clarified by centrifugation at 14,000 rpm for 10 minutes. Thirty μg of protein was electrophoresed in 7.5% polyacrylamide gels, transferred to PVDF membranes for immunoblot analysis with mouse anti-BAF47 antibody (cat. 612110, BD Biosciences, San Jose, CA) and mouse anti-β-actin antibody (cat. A1978, Sigma Aldrich), and developed by chemiluminescence (cat. 170-5061, BioRad Clarity Western ECL Substrate, Hercules, CA) per the manufacturer’s protocol.

### Cell Lines

The VA-ES-BJ cell line was purchased commercially (cat# CRL-2138, ATCC, Manassas, VA). The ESX cell line was provided by author TT^33^. The FU-EPS-1 cell line was provided by author JN^32^.

### Patient Derived Xenograft (PDX) model development

All aspects of cancer tissue sharing for model development were reviewed and approved by the Oregon Health & Science University Institutional Review Board. The PCB490 PDX model was generated at JAX (model number J00007860) by implanting surgical human tumor tissue into 4-6-week-old female immunodeficient NOD.Cg-*Prkdc*^*scid*^ *Il2rg*^*tm1Wjl*^/SzJ (NSG) mice without prior *in vitro* culturing of the tumor cells. Time from surgery to implantation was approximately 24 hours. Once a xenografted tumor reached ∼1000 mm^3^, the tumor was harvested and divided into 3-5 mm^3^ fragments. Fragments were implanted into five 6-8-week-old female NSG mice for expansion to P1. Other fragments were sent for quality control assessment (see below). The remaining fragments were cryopreserved in 10% DMSO. When P1 tumors reached 1000mm^3^ they were harvested and divided into quarters: ¼ for quality control, ¼ snap frozen for genomics, ¼ placed into RNALater (Ambion) for RNA-seq, and the remaining ¼ divided into 3-5 mm^3^ pieces and cryopreserved in 10% DMSO.

The quality control procedures employed for PDX model development included testing the patient tumor for LCMV (lymphocytic choriomeningitis virus), bacterial contamination, and tumor cell content. The engrafted tumors at P0 and P1 were DNA fingerprinted using a Short Tandem Repeat (STR) assay to ensure model provenance in subsequent passages.

Model details available online at: http://tumor.informatics.jax.org/mtbwi/pdxDetails.do?modelID=J000078604

Immunohistochemistry (IHC) for human CD45 (IR75161-2, Agilent Technologies) was performed on paraffin embedded blocks of engrafted tumors to identify cases of lymphomagenesis which have been reported previously in PDXs. IHC for human ki67 (IR62661-2, Agilent Technologies) was used to ensure the propagated tumors were human in origin. H&E sections of engrafted tumors were reviewed by a board-certified pathologist (RGE) to evaluate concordance of the morphological features of the engrafted tumor to the patient tumor. Further, tissue was stained with vimentin (IR63061-2, Agilent Technologies) to confirm human origin.

Model information is publicly accessible at: http://tumor.informatics.jax.org/mtbwi/pdxSearch.do

### Chemical screens

Four chemical screens were used to generate functional drug screening data. The first screen was a custom 60 agent chemical screen of well-characterized target inhibitors denoted the Pediatric Preclinical Testing Initiative Screen Version 2.1 (PPTI screen). Chemical concentrations of agents in all chemical screens were either [10nM, 100nM, 1μM, 10μM] or [100nM, 1μM, 10μM, 100μM] depending on compound activity range. Fifty-four of the 60 drugs on the chemical screen have a published quantified drug-target inhibition profile.

The second screen was a custom 60 agent chemical screen denoted Drug Screen V3 consisting of a variety of small molecule kinase inhibitors, epigenetic target inhibitors, and cell cycle inhibitors. Fifty-two of 60 drugs on the chemical screen have a published drug-target inhibition profile.

The third chemical screen was a GlaxoSmithKline open access Orphan Kinome-focused chemical screen (denoted GSK screen) consisting of 402 novel and newly characterized compounds^39^ with target inhibition profiles quantified by Nanosyn Screening and Profiling Services. Drug-target interaction was assayed over 300 protein targets for each of the 402 compounds. The compounds were tested at 100 nM and 10μM concentrations to bracket the drug-target EC_50_ values. The final EC_50_ values used for analysis of the chemical screen results were inferred from the available data using hill curve fitting to predict the 50% inhibition point.

The final screen was a Roche-developed open access chemical screen (denoted Roche screen) consisting of 223 novel kinase inhibitor compounds^40^. Roche screen compounds had a mixture of quantified and qualified drug-target inhibition profiles, though drug-target inhibition profiles were made available only for sensitive compounds.

Cell cultures were plated in 384-well plates at a seeding density of 5000 cells per well onto gradated concentrations of drug screen compounds. Cells were incubated in model-specific culture media at 37°C, with 5% CO_2_, for 72 hours. Cell viability was assessed by CellTiter-Glo^®^ Luminescent Cell Viability Assay (cat. G7570, Promega, Madison, WI) per manufacturer’s protocol. Luminescence was measured using a BioTek Synergy HT plate reader (BioTek, Winooski, VT). Single agent IC_50_ values were determined using a hill curve-fitting algorithm with variable hill slope coefficients performed in Microsoft Excel. Manual curation and re-fitting of the results was performed before results were finalized.

U23674 primary tumor culture was assayed via three drug screens: PPTI drug screen, GSK drug screen, and the Roche drug screen (Supplemental Figs. S1-3, Supplemental Tables S1-7).

S1-12 primary tumor culture was screened using the PPTI screen (Supplemental Table S21). PCB197 primary tumor culture was screened using the PPTI screen (Supplemental Table S21). PCB490-3, PCB490-4, PCB490-5 primary cultures were screened with Drug Screen V3 and the Roche drug screen (Fig. 3, Supplemental Tables S15-16). Cell lines ESX, FU-EPS-1, and VA-ES-BJ were screened with Drug Screen V3 (Supplemental Table S20). PCB495 primary culture was screened with Drug Screen V3 (Supplemental Table S20).

### U23674 drug combination studies and calculation of Combination Index (CI)

U23674 drug combination validation experiments were guided by GlaxoSmithKline chemical screen PTIM models. Single agent validations to calculate independent drug efficacy were performed at dosages in the range of 5 nM to 100 μM to bracket IC_50_ and IC_25_ dosage values; for combination experiments, the IC_25_ dosage for one agent was tested in combination with gradated dosages (5 nM to 100 μM) of the complementary agent, and vice versa. Single agent and combination agent validation experiments were performed at passage 5.

CI values were generated using the CompuSyn software tool. Effect values for CompuSyn monotherapy and combination therapy were determined by mean cell death based on n = 3 independent experiments with n = 4 technical replicates for the following treatment options: OSI-906, GDC-0941, OSI-906 + GDC-0941 (OSI-906 at IC_25_ + GDC-0941 at varying dosage, OSI-906 at varying dosage + GDC-0941 at IC_25_). CompuSyn CI values were calculated using the non-constant combination setting^41^ (Supplemental Table S13).

We performed low-dose validation experiments to verify PTIM-identified synergistic mechanisms of action; reduced dosages of the combination agents were set to 5 times the EC_50_ value for the predicted target (175 nM OSI-906, 50 nM GDC-0941). CompuSyn CI values to validate the mechanism of synergy were calculated using the non-constant combination setting^41^ (Supplemental Table S13).

In both regular dose and low dose experiments, CI values are reported only for functionally relevant dosages, *i.e.* dosages between the drug target’s EC_50_ and the drug’s maximum achievable human clinical dosage (C_max_). For OSI-906, the functional range is approximately [10 nM, 5 μM] (mouse pharmacokinetics: ∼16 μM C_max_, 6.16 μM C_ss_; human pharmacokinetics: ∼1.481 μM C_max_, 720 nM C_ss_). For GDC-0941, the functional range is approximately [5 nM, 1 μM] (mouse pharmacokinetics: ∼12 μM C_max_, 1.59 μM C_ss_, human pharmacokinetics: ∼1.481 μM C_max_, 720 nM C_ss_). CI values outside these ranges are denoted as N/A in Supplemental Table S13.

### U23674 exome sequencing analysis

Somatic point mutations were identified using the Genome Analysis Toolkit^42^ (GATK, version 3.5.0) from the Broad Institute. Captured DNA libraries were sequenced with the Illumina HiSeq 1000 in paired-end mode. The reads that passed the Illumina BaseCall chastity filter were used for subsequent analysis. The mate pairs were pooled and mapped as single reads to the NCBI GRCm38/mm10 reference genome using the Burrows-Wheeler Aligner^43^ (version 0.7.12), with shorter split hits marked as secondary to ensure compatibility with downstream tools. Identified PCR duplicates, defined as reads likely originating from the same original DNA fragments, were removed using Picard Tools MarkDuplicates (version 1.133). Mapping artifacts introduced during initial mapping are realigned using the GATK IndelRealigner, and base quality score recalibration to empirically adjust quality scores for variant calling was performed by the GATK BaseRecalibrator tool. The same process was used to process both the tumor sample and the matched normal tail sample. Variant discovery was performed by MuTect2^44^, with the NCBI GRCm38/mm10 dbSNP database used to filter known polymorphisms present in the paired sample. Variant annotation and effect prediction was performed using SnpEff^45^ using the GRCm38.81 database. Only medium and high impact effect variants are considered for the purpose of downstream analysis and reporting in figures. Exome analysis protocol is based on the GATK Best Practices protocol.

VarScan2 was used for copy number variation analysis of the paired tumor-normal data^46^. The Burrows-Wheeler Aligner was used to align the tumor and normal samples to NCBI GRCm38/mm10 reference genome as described previously. Samtools (version 0.1.19) mpileup tool with minimum mapping quality of 10 was used to generate the pileup file required by the VarScan2 copycaller function; log_2_ exon coverage ratio data from copycaller was segmented using DNAcopy with the undo.splits = “sdundo” parameter, and deviation from the null hypothesis set above 3 standard deviations. Genes in segments with segment mean above 0.25 or below −0.25 and with p-value below 1e-10 were called as gained or lost, respectively. Copy number variation analysis protocol was partly based on the VarScan2 user manual^47^.

### U23674 RNA deep sequencing analysis

RNA sequencing was performed on a low-passage U23674 culture, and on the control sample consisting of regenerating mouse muscle tissue following cardiotoxin injury *in vivo*. The paired-end raw reads were aligned to the NCBI GRCm38/mm10 reference mouse genome using TopHat version 2.0.9^48^ using Bowtie2 as the short-read aligner. Up to two alignment mismatches were permitted before a read alignment was discarded. The aligned reads were assembled into transcripts using Cufflinks version 2.1.1^49^. Differential gene expression of tumor sample vs. control was performed by Cuffdiff using standard parameters. RNA analysis protocol was largely based on the approach described in the Tophat2 publication^50^. Results are provided in Supplemental Table S9.

### PCB490 exome sequencing analysis

Somatic point mutations were identified using the Genome Analysis Toolkit^42^ (GATK, version 3.8.0) from the Broad Institute. Captured DNA libraries were sequenced in paired-end mode via the BGISeq 500 system at Beijing Genomics Institute. The reads that passed the Illumina BaseCall chastity filter were used for subsequent analysis. The mate pairs were pooled and mapped as single reads to the NCBI GRCm38/mm10 reference genome using the Burrows-Wheeler Aligner^43^ (version 0.7.12), with shorter split hits marked as secondary to ensure compatibility with downstream tools. Identified PCR duplicates, defined as reads likely originating from the same original DNA fragments, were removed using Picard Tools MarkDuplicates (version 1.133). Mapping artifacts introduced during initial mapping are realigned using the GATK IndelRealigner, and base quality score recalibration to empirically adjust quality scores for variant calling was performed by the GATK BaseRecalibrator tool. The same process was used to process both the tumor sample and the matched normal sample. Variant discovery was performed by MuTect2^44^, with the NCBI GRCh38 dbSNP database used to filter known polymorphisms present in the paired sample. Variant annotation and effect prediction was performed using SnpEff^45^ using the GRCh38.87 database. Only medium and high impact variants are considered for the purpose of downstream analysis and reporting in figures. Exome analysis protocol is based on the GATK Best Practices protocol.

VarScan2 was used for copy number variation analysis of the paired tumor-normal data^46^. The Burrows-Wheeler Aligner was used to align the tumor and normal samples to NCBI GRCh38 reference genome as described previously. Samtools (version 1.6) mpileup tool with minimum mapping quality of 10 was used to generate the pileup file required by the VarScan2 copycaller function; log_2_ exon coverage ratio data from copycaller was segmented using DNAcopy with the undo.splits = “sdundo” parameter, and deviation from the null hypothesis set above 3 standard deviations. Genes in segments with segment mean 2 standard deviations above or below ±0.5 and with p-value below 1e-10 were called as gained or lost, respectively. Copy number variation analysis protocol was partly based on the VarScan2 user manual^47^.

### PCB490 RNA deep sequencing analysis

The PCB490 transcriptome library was sequenced with the Illumina HiSeq 2500 in paired-end mode. The reads that passed the chastity filter of Illumina BaseCall software were used for subsequent analysis. The paired-end raw reads for each RNA-seq sample were aligned to the UCSC hg38 reference human genome using Bowtie2 as the short-read aligner^48^ using, allowing up two alignment mismatches before a read alignment was discarded. The aligned reads were assembled into transcripts using Cufflinks version 2.1.1^49^ and quantification was performed with Cuffquant^49^. RNA analysis protocol was adapted from the approach described in the original TopHat2 publication^50^ (Supplemental Table S18).

### RAPID siRNA screen of U23674

U23674 underwent functional single gene knockdown (siRNA interference screen, Supplemental Table S10), however siRNA results were inconsistent with drug screening data (Supplemental Table S11) and are thus relegated to the supplement.

To assess the contribution of individual receptor tyrosine kinases to survival of U23674, we performed RAPID siRNA knockdown screening of U23674. Efficacy of single target knockdown of 85 members of the mouse tyrosine kinase family was performed as previously described^22^. Target sensitivity was determined by resulting cell viability quantified using an MTT assay (M6494; Thermo Fisher Scientific, Waltham, MA, USA). Targets with viability two standard deviations below the mean were identified as high-importance targets^22^ (Supplemental Table S10).

### Phosphoproteomic screen of U23674

U23674 underwent phosphoproteome quantification (Kinexus phosphoproteomics analysis, Supplemental Table S12), however phosphoproteomics results were inconsistent among sample replicates and are thus relegated to the supplement.

To identify differentially phosphorylated protein targets, phosphoproteomics assays (Kinexus, Vancouver, British Columbia, Canada) were used to compare two duplicate cell lysates from U23674 against two duplicate cell lysates from regenerating muscle tissue acting as normal control. To perform the phosphoproteomics analyses, 50 μg of protein lysate from each sample was covalently labeled with a proprietary fluorescent dye. Free dye molecules were removed by gel filtration. After blocking nonspecific binding sites on the array, an incubation chamber was mounted onto the microarray to permit the loading of related samples side by side on the same chip. Following sample incubation, unbound proteins were washed away. Each array produces a pair of 16-bit images, which are captured with a Perkin-Elmer ScanArray Reader laser array scanner. Signal quantification was performed with *ImaGene 8.0* from BioDiscovery with predetermined settings for spot segmentation and background correction. The background-corrected raw intensity data are logarithmically transformed. Z scores are calculated by subtracting the overall average intensity of all spots within a sample from the raw intensity for each spot, and dividing it by the standard deviations (SD) of all of the measured intensities within each sample (Supplemental Table S11).

### Probabilistic Target Inhibition Maps

The Probabilistic Target Inhibition Map (PTIM) approach considers that the underlying mechanism for sensitivity to targeted drugs can be represented by a combination of parallel target groups (all parallel targets need to be inhibited to slow or stop tumor proliferation, similar to Boolean ‘AND’ logic) and series target groups (inhibiting any all targets in any target group will slow or stop tumor proliferation, similar to Boolean ‘OR’ logic). For estimating the series and parallel targets, we analyze cancer cell response to multi-target single agent drugs with overlapping but distinct target sets. For instance, drugs having the same selective target (such as pelitinib and erlotinib, which are potent inhibitors of the kinase target *EGFR*) can show different sensitivity *in vitro* which can be attributed to the biologically relevant side targets of the drugs. Our framework considers primary and secondary drug targets and generates logical groupings of targets (as single-target or multi-target blocks) that best explain chemical screen response data. We now incorporate secondary information to refine PTIM models.

#### PTIM circuit models

PTIM models are visually represented as circuit models. Each “block” in the circuit represents a combination of two or more targets that explain sensitivity of a set of single agent compounds from drug screens. The drug set represented by an individual block is determined by the PTIM objective function and feature selection algorithm^14,16^, and depends on the biological data inputs to the PTIM algorithm.

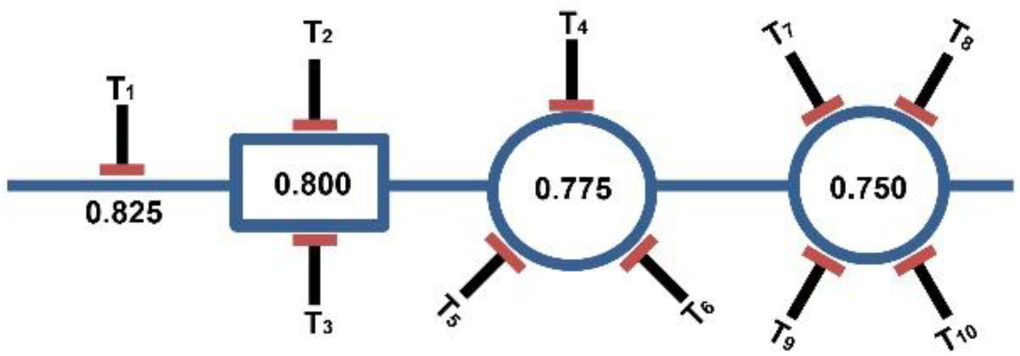

PTIM circuits consist of multiple blocks. Generally, only target combinations of one to four targets are considered during PTIM modeling. Blocks of one target (represented as single inhibitor symbol, T_1_) are called “single points of failure”, i.e. single targets which alone explain the sensitivity of one or more drug screen agents. Combinations of two targets are visually represented by a rectangular block with two inhibitor symbols (block T_2_ – T_3_). Combinations of three targets are visually represented by a circular block with three inhibitor symbols (block T_4_ – T_5_ – T_6_). Combinations of four targets are visually represented by a circular block with four inhibitor symbols (block T_7_ – T_8_ – T_9_ – T_10_). Each block has an associated score value (e.g. 0.825, 0.800, 0.775, 0.750, respectively) that represents the scaled sensitivity of all drug screen agents grouped in the block’s target combination^14,16^. In brief, all single agent sensitivities (as IC_50_ values) are log_10_ scaled and converted to [0,1] sensitivity values via the following equation:

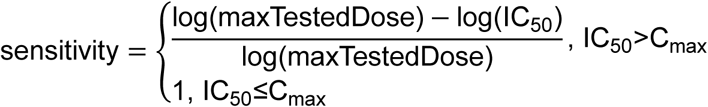

Thus, the lower the IC_50_, the higher the sensitivity score. The score assigned to each block is a determined by the sensitivity of the drug screen agents assigned to the block following several correction factors^14,16^. The shape of blocks in PTIM circuits are meant to serve as a convenient visual representation; ordering of PTIM circuit blocks are determined by overall score, with highest scored blocks on the left descending to lowest scored blocks on the right. The general PTIM algorithm is presented in previously published work^14,16–18^. Methods for integration of secondary biological data are provided in the methods sections for modeling of U23674 and modeling of PCB490.

#### Synergy, heterogeneity, and resistance via PTIM models

PTIM circuit models are also designed to visually represent the clinical challenges PTIM modeling seeks to address. Synergistic drug combinations can be selected for any block with two or more targets by selecting two (or more) drugs which inhibit all targets in the block; the selected combination should kill cancer cells while monotherapy treatment would not. For example, based on (block T_2_ – T_3_), a drug inhibiting T_2_ and a drug inhibiting T_3_ will individually not slow tumor growth for the sample patient, while the combination T_2_ + T_3_ will.

Drug screening multiple spatially-distinct sites from a solid tumor can result in heterogeneous single agent sensitivity. Target group blocks identified as common amongst PTIM models from each distinct region can be used to design a drug combination that should slow or stop tumor growth across the entire heterogeneous tumor. Multi-site PTIM models can thus define heterogeneity-aware drug combinations. Each block in a PTIM circuit represents a set of effective treatment options; effective options on parallel biological pathways represent multiple distinct treatment options which can individually slow tumor growth. A drug combination which inhibits multiple parallel biological pathway blocks can shut down potential survival mechanisms for cancer cells, thus abrogating development of resistance. Series PTIM blocks can thus define resistance abrogating drug combinations.

### Integrative Nonlinear Boolean Modeling for U23674

Probabilistic Target Inhibition Maps (PTIMs) were used for integrative analysis of U23674 biological data^16–18^.

#### RNA-seq integration

For targets common to both RNA expression data and drug-target interaction data, we use gene expression data to eliminate possible false positives from chemical screen results and to narrow down the true positives among relevant targets identified by the PTIM approach. False positives are identified as targets that are inhibited by effective drugs but are not expressed in cancer cells at levels above matched normal cells. Note that we consider the effect of a molecularly-targeted drug is to inhibit the target when it is expressed, thus under-expressed drug targets will have limited impact on drug response. Here, over-expression is determined as gene expression in the tumor sample 50% greater than that in the control sample. The RNA-seq target set is used for PTIM creation via the published model development algorithms.

Formally, RNA-seq data is integrated as below:

- T ≔ targets inhibited in drug screen
- G ≔ targets with RNA-seq expression in tumor and normal cells
- Tumor(x) ≔ gene expression of target x in tumor sample
- Normal(x) ≔ gene expression of target x in normal sample
- ∀_x ∈ T∩G_ Ratio(x)≔ Tumor(x) Normal(x)
- 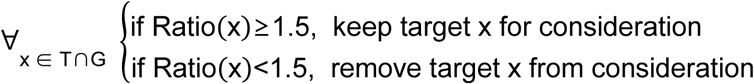
- ∀_x ∉ T∩G_ keep target x for consideration

#### Exome-seq integration

We use exome sequencing data to identify targets likely important in the biological function of tumor cells. We assume that variations may explain the behavior of compounds inhibiting the mutated/altered targets. Depending on the available evidence for mutations and variations, targets are incorporated into the model search or final PTIM model via the published model development algorithms.

Formally, exome-seq data is integrated as below:

- T ≔ targets inhibited in drug screen
- G ≔ targets with RNA-seq expression in tumor and normal cells
- Mut(x) ≔ mutation/indel status of target x (low/med/high impact mutation/indel)
- CNV(x) ≔ copy number status of target x (gain/loss)
- 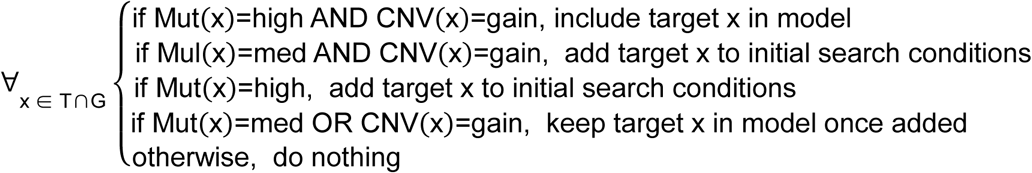
- ∀_x ∉ T∩G_ keep target x for consideration

#### RAPID siRNA screen integration

RAPID screen results identify high sensitivity single target mechanisms of cancer cell growth inhibition; identified hit targets were set as “required” (forced inclusion) in the RAPID siRNA PTIM model effective as sensitive siRNA targets may explain drug sensitivity of agents inhibiting the siRNA targets. Targets not identified by RAPID screening could still have effect in multi-target combinations, and thus were retained for consideration. The RAPID target set is used for PTIM creation via the published model development algorithms.

Formally, RAPID siRNA data is integrated as below:

- T ≔ targets inhibited in drug screen
- G ≔ targets with RAPID siRNA viability data
- RAPID(x) ≔ cell viability following siRNA knockdown of target x
- (μ, σ) ≔ mean and standard deviation of RAPID siRNA dataset
- 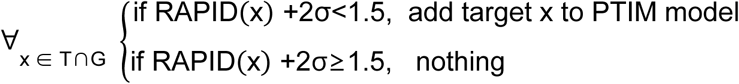
- ∀_x ∉ T∩G_ keep target x for consideration

#### Kinexus phosphoproteomics screen integration

the phosphoproteomics screen results identify differentially phosphorylated targets and associated pathways, phosphorylation of these targets may be pushing the system towards a particular phenotype, and intervention in the form of changing phosphorylation status might result in significant changes to the system. Targets identified as overactive in tumor compared to normal are included in the target set for the PTIM model. The phosphoproteomics target set is used for PTIM creation via the published model development algorithms.

- T ≔ targets inhibited in drug screen
- G ≔ targets with RAPID siRNA viability data
- P_1_(x) ≔ z-score ratio of target x in U23674 replicate 1 vs normal
- P_2_(x) ≔ z-score ratio of target x in U23674 replicate 2 vs normal
- 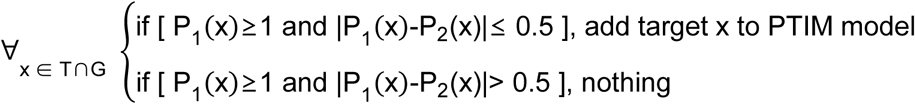
- ∀_x ∉ T∩G_ keep target x for consideration

### Integrative Nonlinear Boolean Modeling for PCB490

Probabilistic Target Inhibition Maps (PTIMs) were used for integrative analysis of heterogeneous PCB490 biological data^16–18^.

### RNA-seq integration

RNA sequencing data for PCB490-5 was used to eliminate under-expressed targets from consideration for PTIM model development, reducing the potential number of models. Due to possessing only tumor tissue for PCB490, RNA sequencing was performed only on the tumor sample; targets with quantified expression above the first quantile were retained for PTIM model development. The RNA-seq target set is used for PTIM creation via the published model development algorithms.

Formally, RNA-seq data is integrated as below:

- T ≔ targets inhibited in drug screen
- G ≔ targets with RNA-seq expression in tumor and normal cells
- Tumor(x) :=gene expression of target x in tumor sample
- Q_1_≔first quartile of Tumor(*) data
- 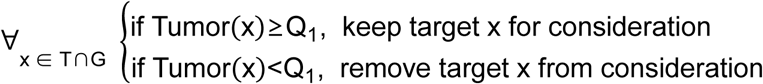
- ∀_x ∉ T∩G_ keep target x for consideration

#### PTIM Ensemble combination optimization

To address tumor heterogeneity concerns, PTIM computational models were generated for each of the drug screened PCB490 cultures (PCB490-3, PCB490-4, and PCB490-5). The PCB490-5 PTIM model integrates RNA sequencing data as above. Combination therapy for PCB490 was designed by identifying PTIM target blocks in each of the three different cell models druggable by the same two-drug combination.

### Rewiring experiments for U23674

Untreated U23674 cells were screened using the Roche Orphan Kinome Screen and concurrently used to establish 6 additional independent cultures grown in culture media at 37°C with 5% CO2. Upon reaching 70% confluence, low dosage single agents and drug combinations (DMSO vehicle, 175 nM OSI-906, 50 nM GDC-0941, 175 nM OSI-906 + 50 nM GDC-0941) were added to culture plates and incubated for 72 hours (Supplemental Fig. S9). Cell plates were then washed in Phosphate Buffered Saline (PBS, Gibco, Grand Island, New York), trypsonized with Trypsin-EDTA (0.25%) (25200056, Thermo Fisher Scientific), and screened using the Roche Orphan Kinome Screen (Supplemental Fig. S10, Supplemental Table S14). Rewiring data was used to generate PTIM models to identify post-intervention changes to U23674 survival pathways (Supplemental Fig. S11).

### Resistance abrogation experiments for PCB197 and S1-12

PCB197 PPTI screen data and S1-12 PPTI screen data were used to generate PTIM models to identify canine and human cross-species mechanistic targets for undifferentiated pleomorphic sarcoma. Consensus targets were chosen for their appearance in human and canine PTIM models; two drugs (obatoclax, an MCL1 inhibitor and panobinostat, a pan-HDAC inhibitor) that most effectively inhibited PTIM-identified blocks at clinically achievable concentrations were selected for validation.

Potential for resistance abrogation by targeting 2 blocks common to both human and canine PTIM models directed a six-arm proof-of-principle experiment to show that inhibition of multiple blocks inhibited could abrogate tumor cell resistance. PCB197 and S1-12 cell cultures were seeded in quadruplicate on 6-well plates (6 plates per cell model) with 10,000 cells per well. Cells were plated 24 hours prior to incubation with any drug. The drug concentrations chosen were 1.5 times the EC_50_ of the PTIM target of interest. The drug selection was based on desired targets, as well as requiring drug concentration for reaching 1.5 times target K_d_ must also be less than the maximum clinically achievable concentration.

One plate per cell model was assigned to each of the 6 treatment arms: 1) vehicle control; 2) obatoclax for 6 days; 3) panobinostat for 6 days; 4) obatoclax for 3 days, wash, then panobinostat for 3 days; 5) panobinostat for 3 days, wash, then obatoclax for 3 days; 6) obatoclax + panobinostat simultaneously for 6 days. After 6 days, culture plates were washed with PBS and fresh DMEM with 10% FBS was placed in each well. Wells were monitored until confluency was observed. Primary study endpoint was days to well confluency as determined by a single user. Cells were also counted manually with a hemocytometer and photographed to confirm consistency of the user’s definition of confluency. If after 100 days the cells did not reach confluency, the cell study ends and the remaining cells are counted. The experimental design and results are available in Fig. 5.

### Orthotopic allograft studies for U23674

We orthotopically engrafted *SHO (*SCID/hairless/outbred) mice (Charles River, Wilmington, Massachusetts) with 10^6^ U23674 cells. Engraftment was performed after injuring the right gastrocnemius muscle by cardiotoxin injection as previously described^22^. Treatment commenced two days after engraftment; mice were treated with vehicle control (tartaric acid + TWEEN80/methylcellulose), 50 mg/kg OSI-906, 150 mg/kg GDC-0941, and combination 50 mg/kg OSI-906 plus 150 mg/kg GDC-0941. Each arm was assigned n = 8 mice per arm. The GDC-0941 arm lost one mouse due to oral gavage; the corresponding data point was censored. Dosing schedule was once daily by oral gavage up to day 5, at which time dosing was performed every other day. The endpoint considered for the study and survival analysis was tumor volume = 1.4cc. All drug studies in mice were performed after receiving approval from the IACUC at Oregon Health and Science University. Sample size was selected to provide 90% power for the statistical tests. Variances between compared groups were similar per Greenwood’s Formula. No blinding was performed during *in vivo* experiments. All animal procedures were conducted in accordance with the Guidelines for the Care and Use of Laboratory Animals and were approved by the Institutional Animal Care and Use Committee at the Oregon Health & Science University.

### Patient Derived Xenograft (PDX) model testing for PCB490

Female stock mice (Envigo *Foxn1*^*nu*^ Athymic nudes) were implanted bilaterally with approximately 5×5×5mm fragments subcutaneously in the left and right flanks with JAX PDX model of Human Epithelioid Sarcoma (J000078604 (PCB490) – JAX-001). After the tumors reached 1-1.5 cm^3^, they were harvested and the viable tumor fragments approximately 5×5×5mm were implanted subcutaneously in the left flank of the female study mice (Envigo *Foxn1*^*nu*^ Athymic nudes). Each animal was implanted with a specific passage lot and documented. J000078604 (PCB490) – JAX-001) was P4. Tumor growth was monitored twice a week using digital calipers and the tumor volume (TV) was calculated using the formula (0.52 × [length × width^2^]). When the TV reached approximately 150-250 mm^3^ animals were matched by tumor size and assigned into control or treatment groups (3/group for J000078604 (PCB490) – JAX-001). Dosing was initiated on Day 0. After the initiation of dosing, animals were weighed using a digital scale and TV was measured twice per week. For J000078604 (PCB490) – JAX-001, sunitinib was administered PO QD for 21 days at 30.0 mg/kg/dose and BEZ235 was administered PO QD for 21 days at 25.0 mg/kg/dose alone and in combination.

### Statistics

Pearson correlation coefficients for PCB490 drug screen response data were calculated in MATLAB using the corrcoef function, correlating drug screen IC_50_ values among all samples. Statistical comparison of correlation coefficients between groups was performed with two-tailed student’s T-test.

The Kaplan-Meier curves for the U23674 *in vivo* orthotropic allograft studies were generated via logrank statistical tests. No blinding was performed.

P-values for the PCB490 PDX experiment were generated using a repeated measures linear model of tumor size in terms of group, time, and the group by time interaction based on an autoregressive order 1 correlation assumption with SAS Version 9.4 for Windows (SAS Institute, Cary, NC).

### Study Approval

All animal procedures performed at Oregon Health & Science University were conducted in accordance with the Guidelines for the Care and Use of Laboratory Animals and were approved by the Institutional Animal Care and Use Committee at the Oregon Health & Science University.

All animal procedures performed at The Jackson Laboratory were conducted in accordance with the Guidelines for the Care and Use of Laboratory Animals and were approved by the Institutional Animal Care and Use Committee at The Jackson Laboratory.

All animal procedures performed at Champions Oncology were conducted in accordance with the Guidelines for the Care and Use of Laboratory Animals and were approved by the Institutional Animal Care and Use Committee at Champions Oncology.

### Data Availability

All analyzed data is available in the supplemental materials for this manuscript. Sequencing data will be made available through the Gene Expression Omnibus (GEO) and the Database of Genotypes and Phenotypes (dbGaP). Accession numbers are not yet available.

## DISCUSSION

The work presented here represents validation experiments for three aspects of PTIM-guided personalized cancer therapy design: drug sensitivity and synergy prediction in a GEMM-origin aRMS, heterogeneity-consensus drug combination design and validation in the first-reported EPS PDX model, and mitigation of cancer cell resistance mechanisms in cross-species *in vitro* validation experiments. Our studies suggest the high value of combining functional screening data with secondary molecular data (especially RNA-seq data) in the design of personalized drug combinations as a supplement to or alternative to DNA sequencing-based therapy assignment. While increased effort is required to generate functional data, the additional information and evidence may prove useful in designing therapeutically effective personalized cancer treatments.

Critically, the timeframe for PTIM-based combination therapy design is less than the time required for standard high-throughput sequencing experiments. The PTIM analysis pipeline can be performed in under 2 weeks and without the explicit need for sequencing results. Currently, the time limiting step in integrative PTIM analysis is exome and RNA sequencing, for which new technology is rapidly reducing time and monetary cost. Functional drug screening in standard well plates can be performed for under $300, and CLIA-certified physical sequencing experiments are now under $500 per analyte per experiment; the cost of a complete functional and molecular analysis now represents a fraction of drug cost and may be accessible to a large population of cancer patients.

These three PTIM-guided validation experiments serve as proofs-of-concept; the critical next stage in PTIM-based personalized therapy design will be prospective evaluation using compassionate use programs in academic institutions, to test therapies as n-of-1 trials in individual patients. As the cost of analysis is low, the major challenges will be 1) administration of FDA-approved drugs, very likely as off-label therapy in combinations potentially not validated in Phase I trials, and 2) financial costs associated with modern targeted therapy regimens, which may currently be prohibitive for some patients.

As drug screen results ultimately guide PTIM modeling, computational modeling of different disease types will require designing disease-specific compound screens to maximize the breadth and depth of disease-relevant multi-target interactions^15^. Similarly, different types of secondary molecular data influences target selection during PTIM model construction depending on the underlying analyte or perturbation, with different secondary datasets expectedly producing different PTIM models. Selection of secondary datasets to generate for individual cases will depend on availability of tumor tissue and expected predictive utility of individual datasets. Based on widespread clinical utility and published studies, the current standard secondary datasets for PTIM modeling are exome sequencing data and RNA sequencing data^14^. As high-throughput analysis of additional biological analytes becomes available through CLIA certified procedures, new datatypes will be integrated into PTIM models. In particular, recent advances in robust generation of proteomics data from patient samples^51–53^ may enable routine integration of proteomics data into PTIM modeling beyond the test case presented in this work.

PTIM-based personalized cancer therapy also requires development of personalized toxicity and dosing prediction methods for designing maximally effective, minimally toxic drug combinations. Research on toxicity prediction is underway, as is research on incorporating chemotherapy backbone into drug combination predictions. While validated PTIM models are currently based on low-passage cell cultures (U23674, PCB490, S1-12, PCB197), future application of PTIM models will use direct-to-plate tumor screening to best recapitulate the patient’s disease state and to remove the dependence on cell culture establishment. Finally, we will pursue expansion of disease-consensus PTIM modeling^19^ to establish new disease-specific drug combinations based on integrated drug screening and high-throughput sequencing data.

PTIM-based personalized combination therapy has been designed to uniquely leverage patient-specific functional and biological data to address some of the critical unmet clinical needs of the 60% of cancer patients for whom tumor DNA analysis is uninformative^2^ and the 600,000 patients lost to cancer every year^1^ who have exhausted clinical options. PTIM modeling can also meet the needs of cancer patients with rare diseases, such as the spectrum of 60+ cancers known as non-rhabdomyosarcoma soft tissue sarcomas (including EPS) for which effective clinical treatments may not exist and may never be developed due to a paucity of disease models for research. These two groups represent a significant cancer patient population for which PTIM modeling may provide evidence-based treatment options where no therapeutic avenues exist.

## Acknowledgements

This work was supported by the Scott Carter Foundation Fellowship grant (to N.E.B), the SuperSam Foundation Fellowship grant (to N.E.B), the Prayers for Elijah Foundation (to C.K. and N.E.B.), NSF award CCF0953366 (to R.P.), the Damon Runyon-Sohn & St. Baldrick’s Foundation training grants (to L.E.D.), the AAO-HNSF Saidee Keller Memorial Resident Research Grant, American Academy of Otolaryngology – Head & Neck Surgery Foundation (AAO-HNSF) and the Centralized Otolaryngology Research Effort (CORE) Study Section (to M.N.G), as well as by a gift from an anonymous donor. We thank William Zuercher and David Drewry at GlaxoSmithKline and Paul Gillespie at Roche for making the respective compound libraries available for the community and this study.

### FINANCIAL DISCLOSURE

Investigators N.E.B., C.K. and R.P. have previously filed invention disclosures for the probabilistic Boolean model that integrates chemical screening and genomics data, and are in the process of forming a related company. The authors have declared these conflicts to their respective institutions, which are developing conflict of interest management plans.

All other authors declare no competing interests.

---

**Supplemental Figure 1.**
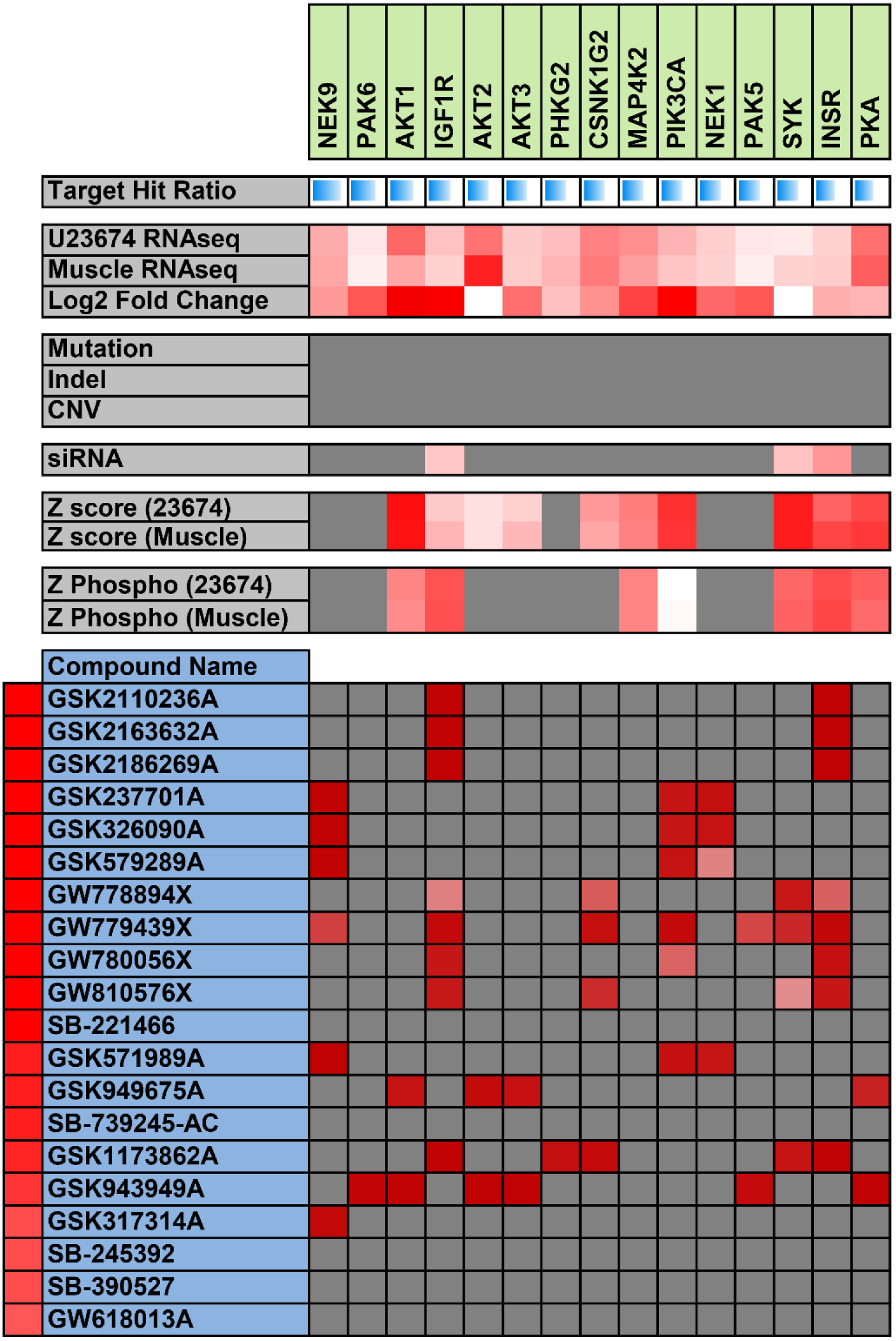
Heat map of merged chemical screen, RNA-seq, siRNA, and phosphoproteomics results for GlaxoSmithKline (GSK) Orphan Kinome screen. Due to the large number of compounds and protein targets, only a limited scope of compounds and targets is shown here (for full data, see Supplemental Table 1). Bright red indicates high sensitivity values, gradating down to white meaning low sensitivity. Gray indicates no interaction or no available data. Asterisk indicates targets later validated *in vivo*.

**Supplemental Figure 2.**
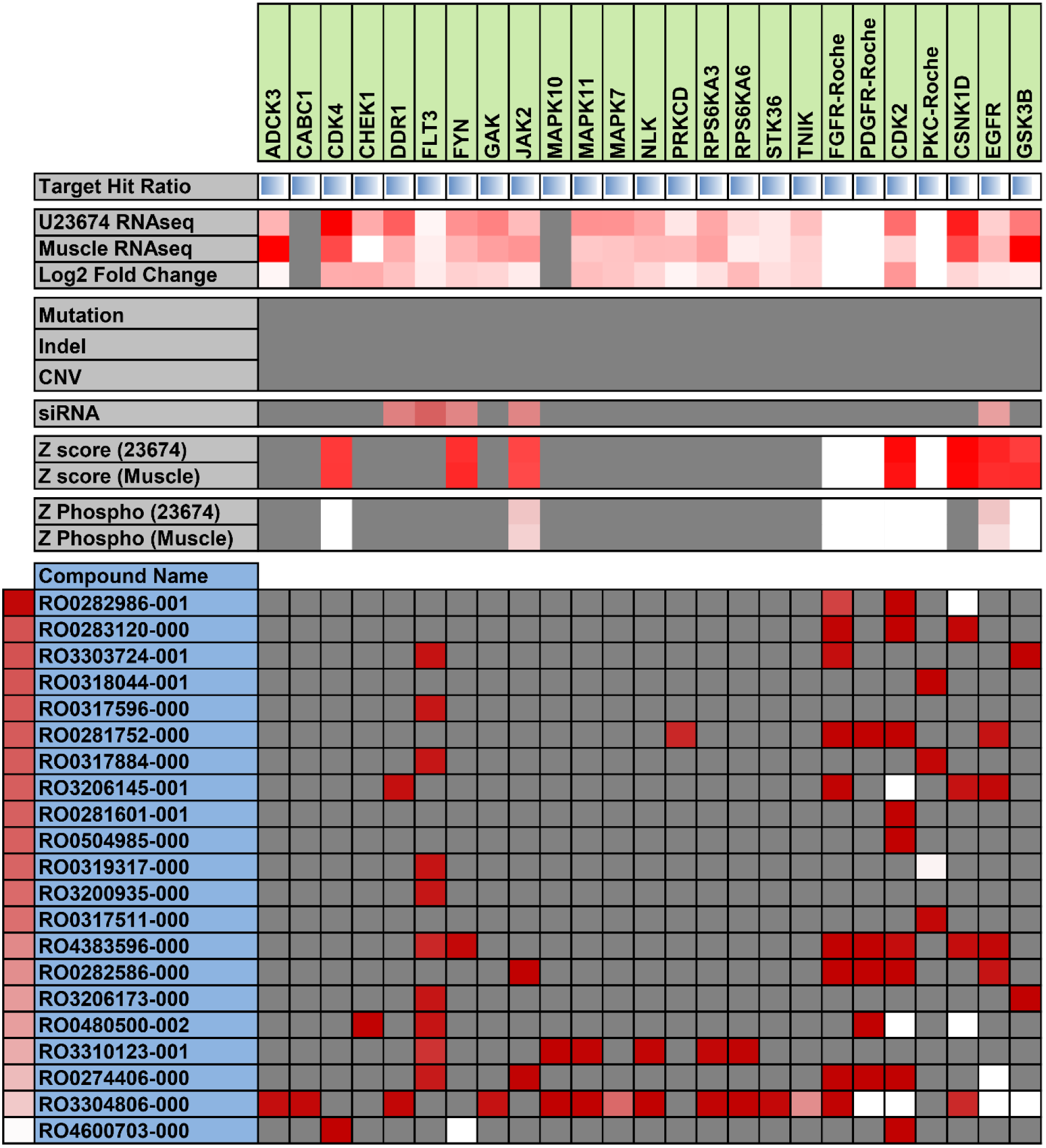
Heat map of merged Roche Orphan Kinome chemical screen, RNA-seq, siRNA, and phosphoproteomics results. Due to the large number of compounds and protein targets, only a limited scope of compounds and targets is shown here (For full data, see Supplemental Table 3). Bright red indicates high sensitivity values, gradating down to white meaning low sensitivity. Gray indicates no interaction or no available data.

**Supplemental Figure 3.**
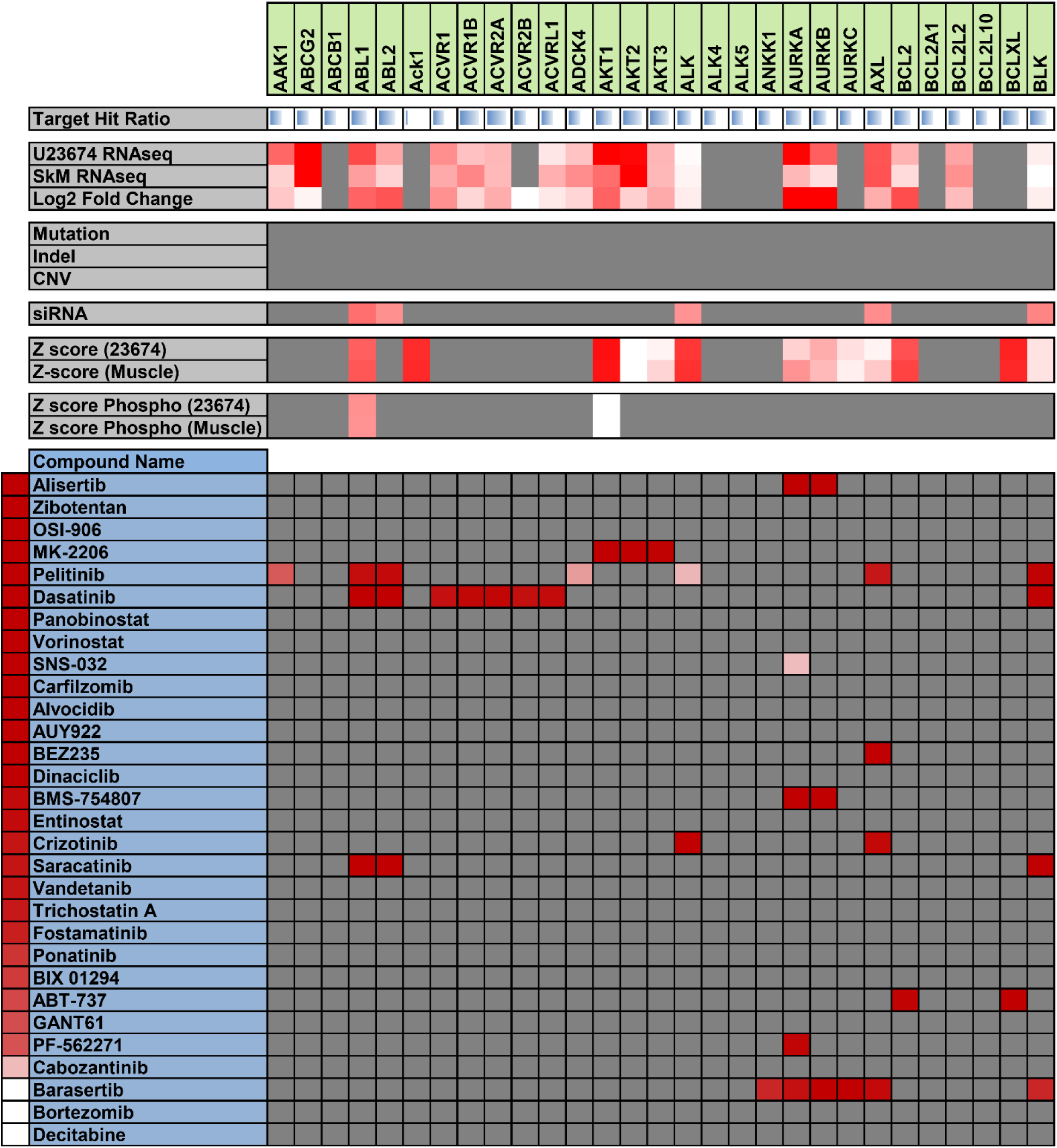
Heat map of joint version 2.1 chemical screen, RNA-seq, siRNA, and phosphoproteomics results. Due to the large number of compounds and protein targets, only a limited scope of compounds and targets is shown here (For full data, see Supplemental Table 7). Bright red indicates high sensitivity values, gradating down to white meaning low sensitivity. Gray indicates no interaction or no available data.

**Supplemental Figure 4.**
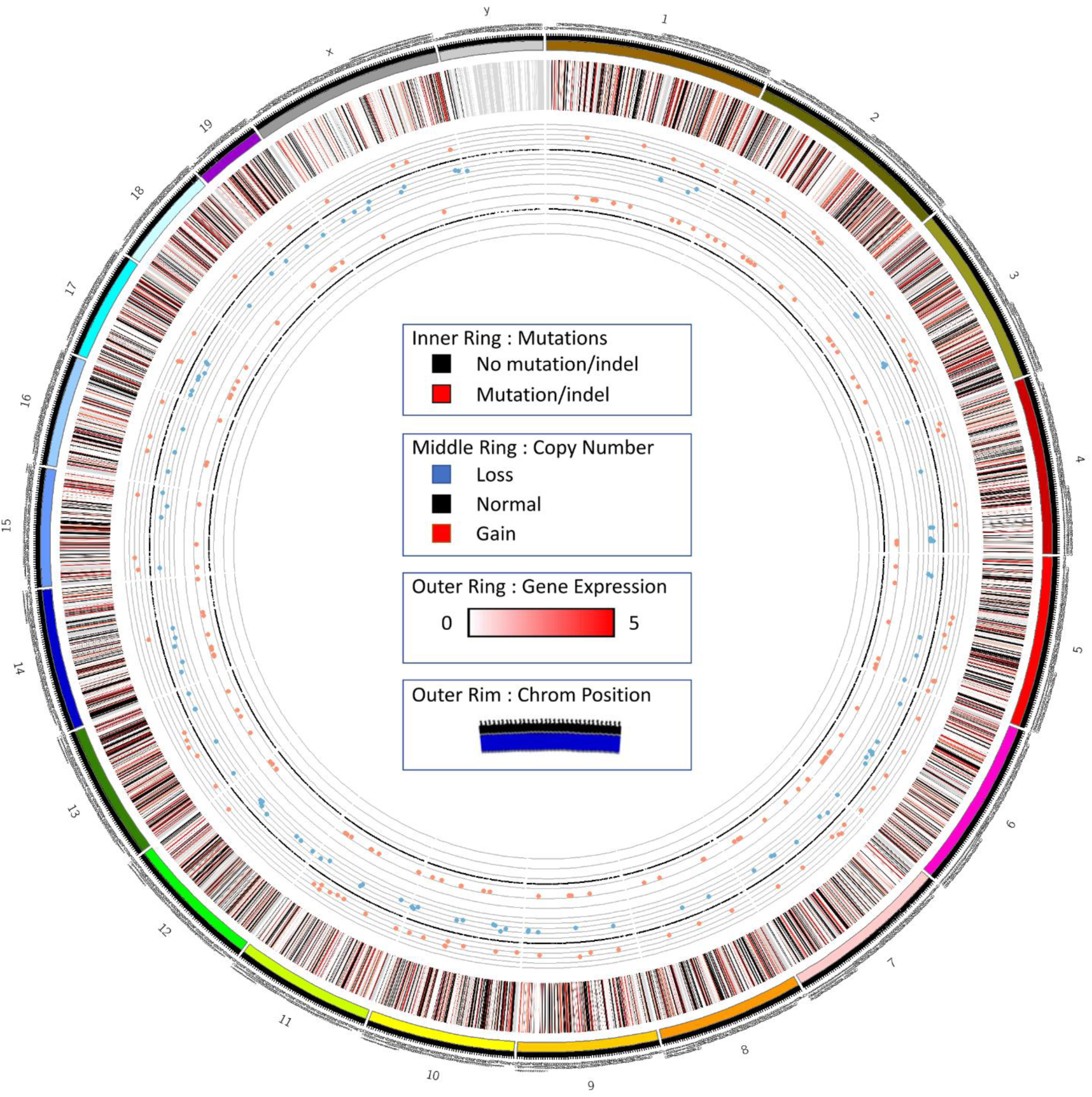
Circos plot of U23674 RNA sequencing and exome sequencing data. The outermost data circle represents log_2_-scaled gene expression [log_2_(expression+1), low expression (white) to high expression (red), with missing values colored black]. The middle circle represents genes with identified mutations or indels (red) or lack thereof (black). The innermost circle represents copy number variations (red is amplification, blue is deletion, black is no variation).

**Supplemental Figure 5.**
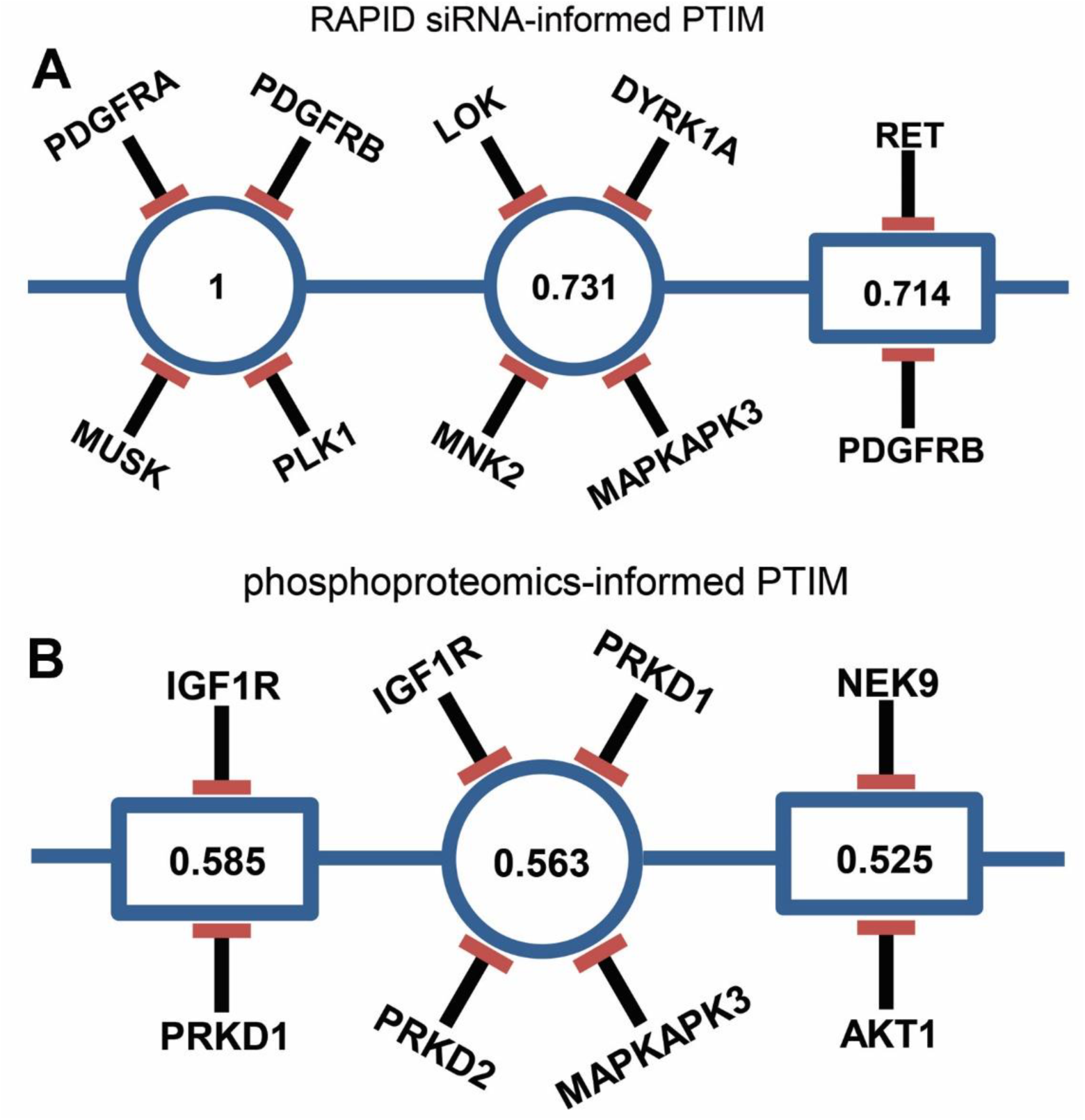
PTIM models developed using secondary datasets. (**A**) siRNA + Chemical screen informed PTIM. Values in the center of PTIM blocks represent expected scaled sensitivity following inhibition of associated block targets. (**B**) phosphoproteomics + Chemical screen informed PTIM. The values within the target blocks indicate scaled drug sensitivity^16^ when block targets are inhibited.

**Supplemental Figure 6.**
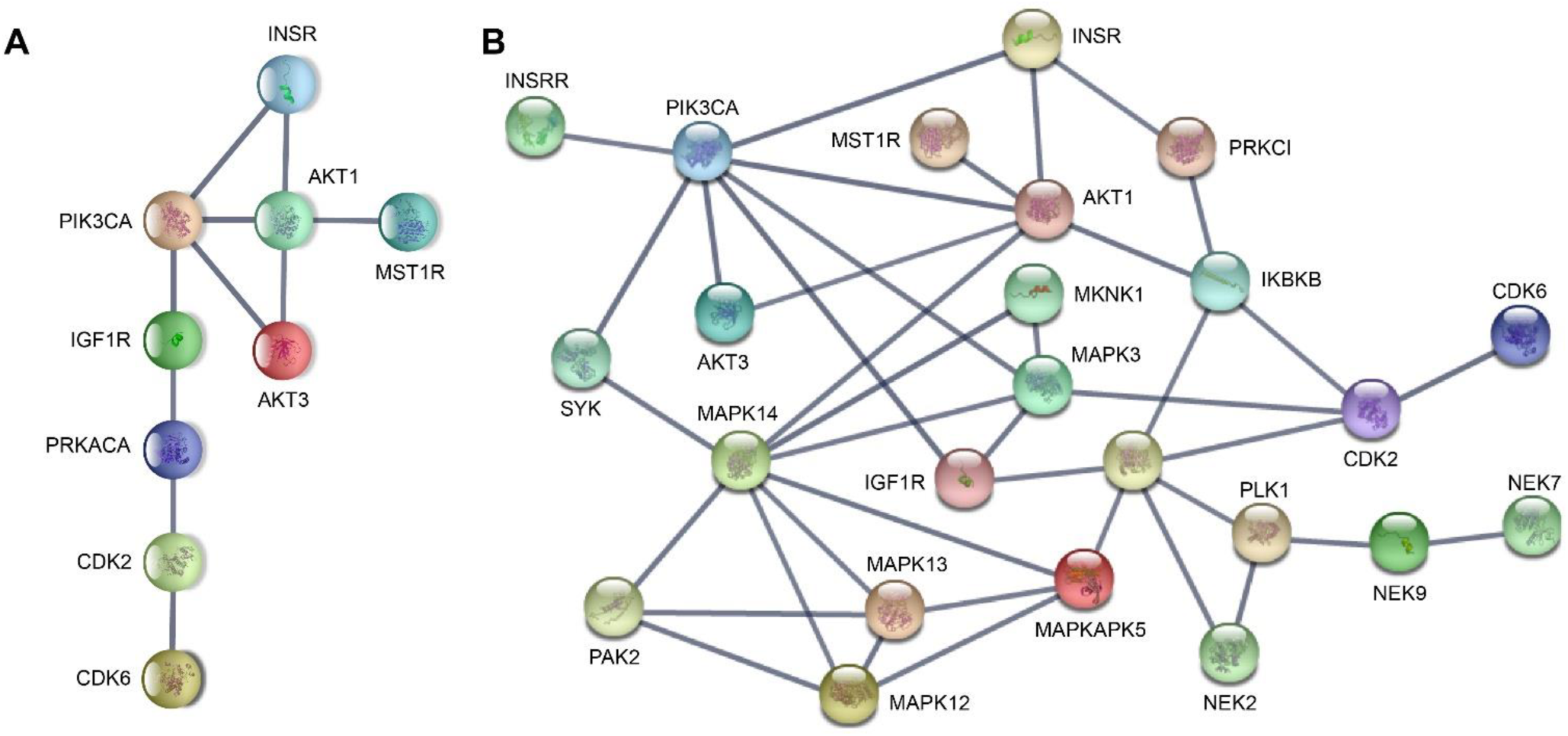
STRINGdb visualizations of protein-protein interaction networks implicated by PTIM models. The protein-protein interaction networks here are derived from targets selected to define drug sensitivity during PTIM modeling. Edges in the STRINGdb graph represent confidence of interactions based on data from multiple published sources. Edges with confidence > 0.9 are represented on the graph. The asterisk indicates targets validated *in vitro.* (**A**) Network of the set of targets common to the models developed for the GSK Orphan Kinome screen and the PPTI screen. Enrichment p-value < 0.01. (**B**) Network of the targets identified by the GSK Orphan Kinome screen alone. Enrichment pvalue < 0.01.

**Supplemental Figure 7.**
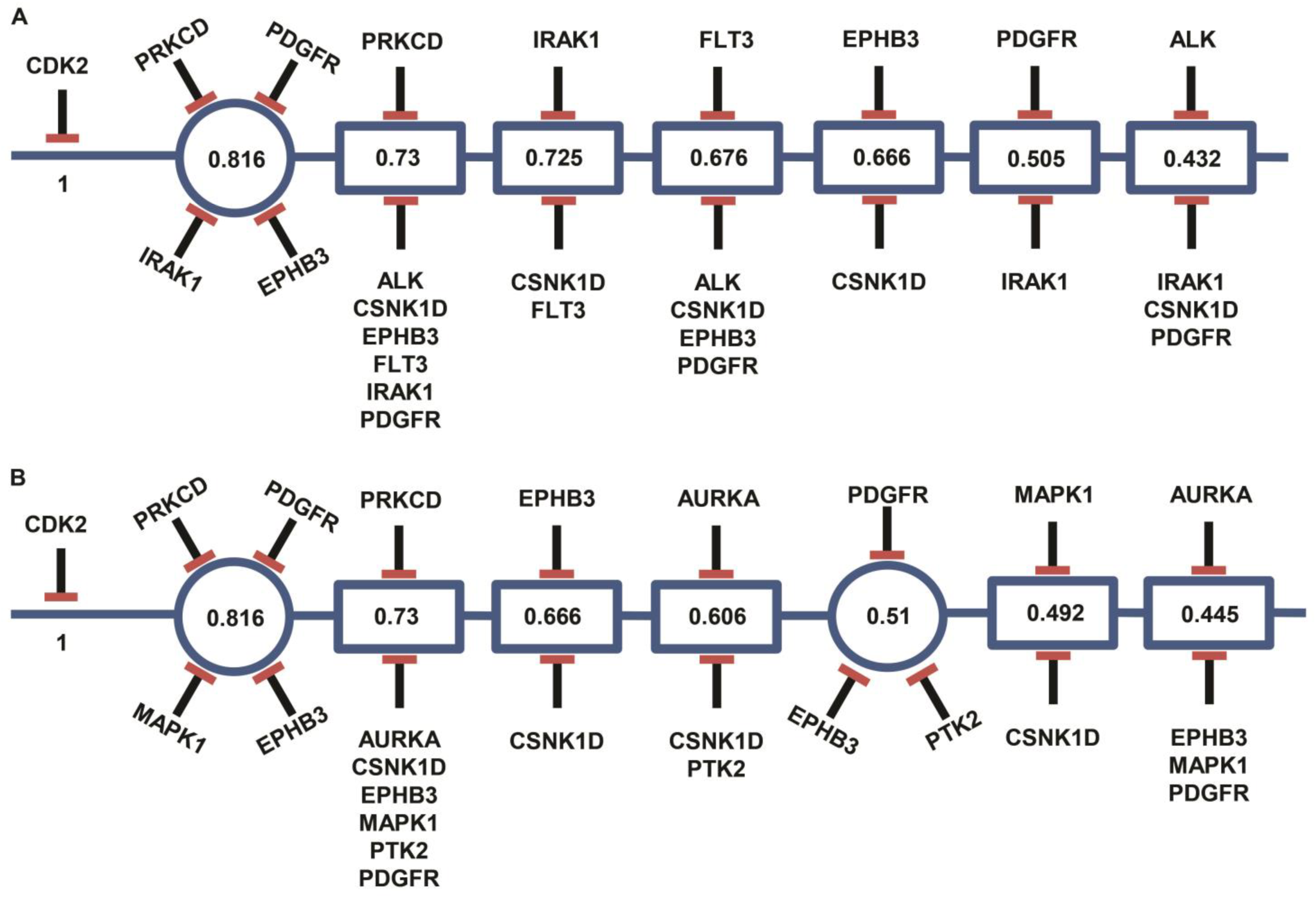
Probabilistic Target Inhibition Map (PTIM) model of U23674 Roche chemical screen hits. Values in the center of PTIM blocks represent expected scaled sensitivity following inhibition of associated block targets. (**A**) Base chemical screen informed PTIM. (**B**) RNA-seq + chemical screen informed PTIM. Roche screen hits include CDK2 inhibitors. However, no CDK inhibitor was a known inhibitor of non-CDK targets, limiting development of personalized combinations involving CDK inhibitors.

**Supplemental Figure 8.**
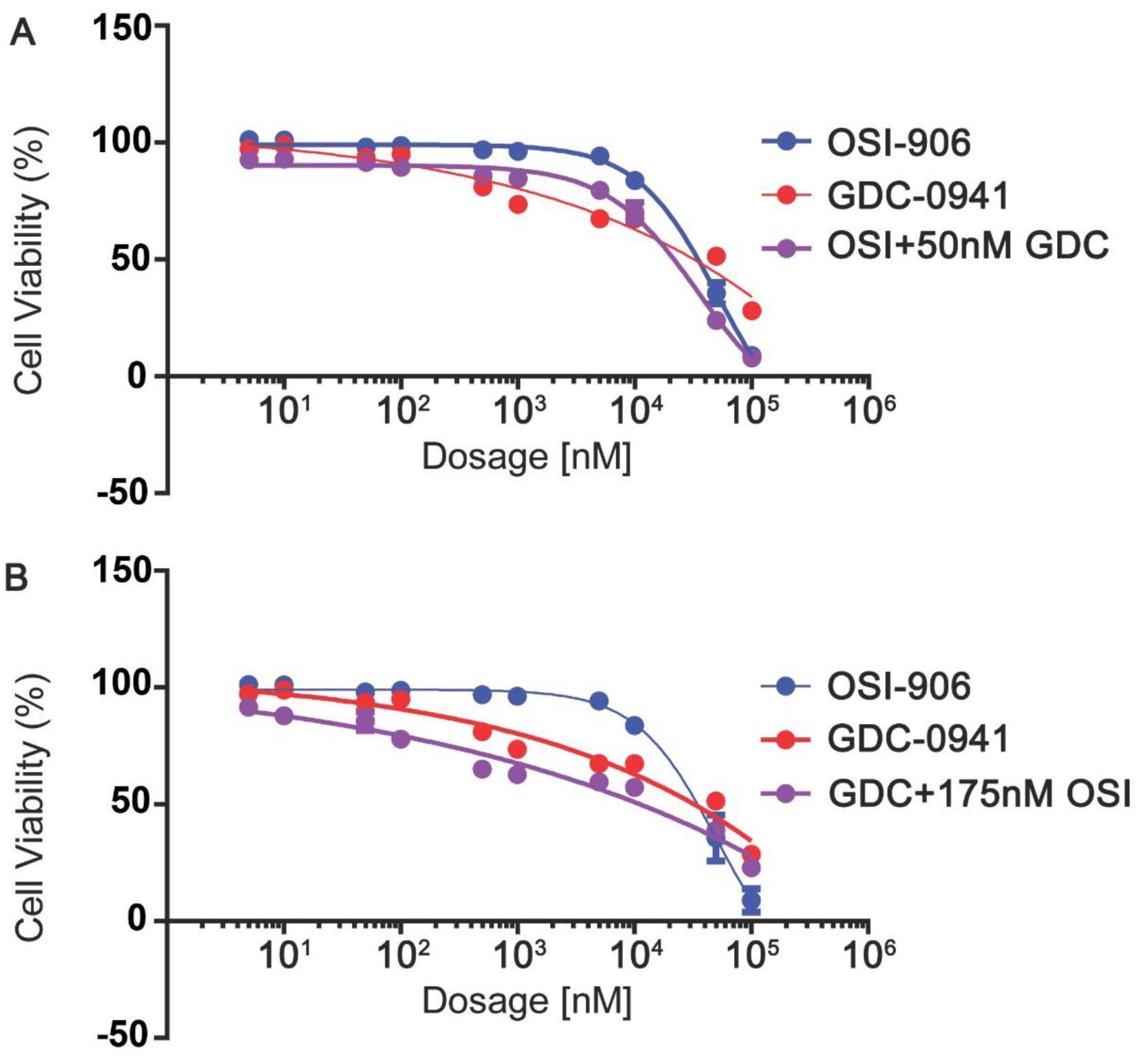
Low dose combination validation results for drug combinations GDC-0941 + OSI-906. Results are based on n = 3 independent experiments with n = 4 replicates. (**A**) Dose response curve for OSI-906 varied dosage + GDC-0941 low fixed dosage. The response for GDC-0941 at varied dosages is included. (**B**) Dose response curve for GDC-0941 varied dosage + OSI-906 low fixed dosage. The response for OSI-906 at varied dosages is included.

**Supplemental Figure 9.**
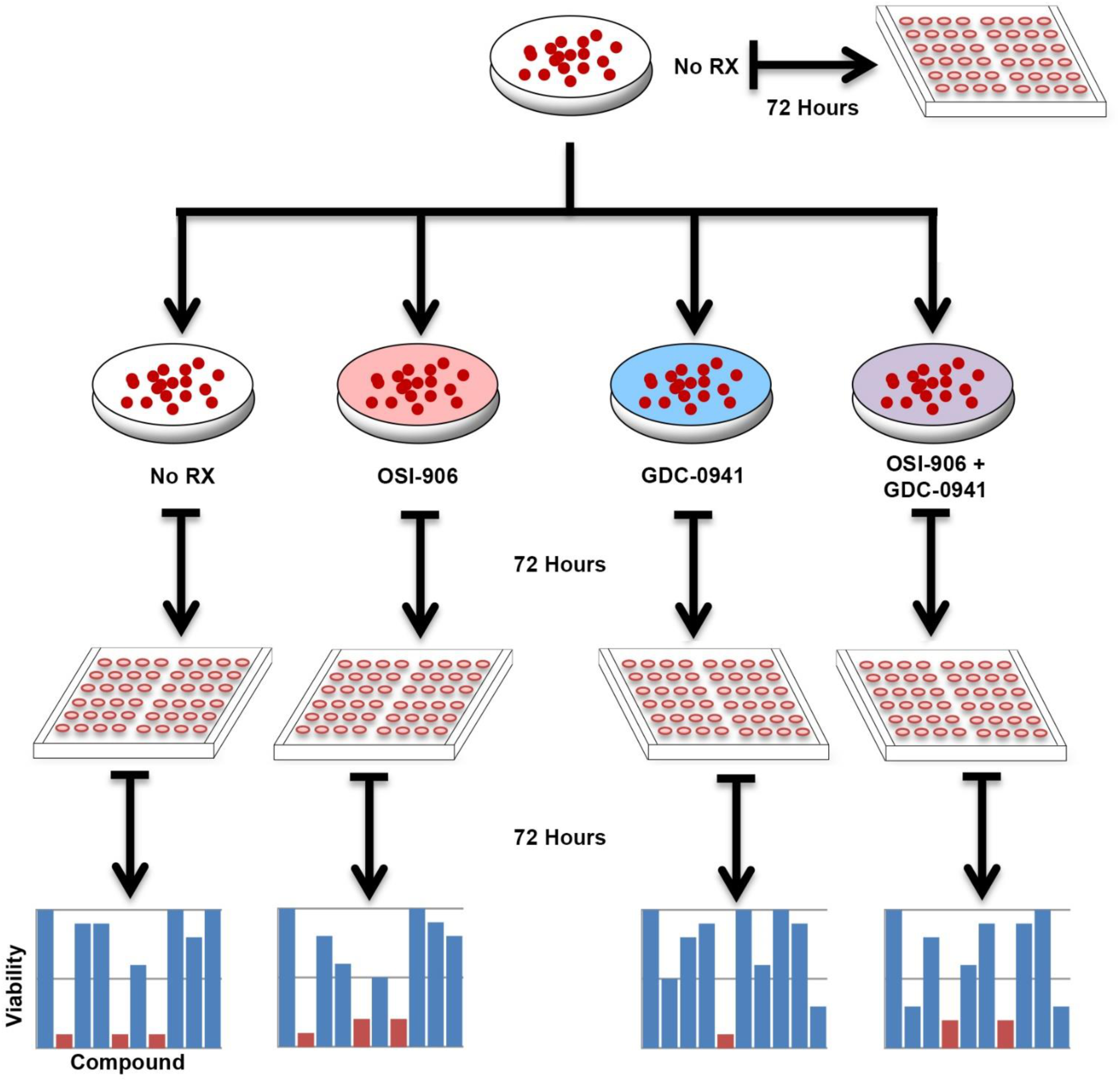
Schematic of PTIM-informed U23674 rewiring experiment. An initial culture of U23674 is screened using the Roche screen. The same culture is used to seed 6 new cultures, which are grown until the cell population is sufficient for drug screening. Five of the 6 cultures were treated using single agents and drug combinations in low dosages (75 nM OSI-906, 50 nM GDC-0941) and one culture was left untreated. After treatment and incubation for 72 hours, the compounds were removed the cells were screened using the Roche Orphan Kinome screen.

**Supplemental Figure 10.**
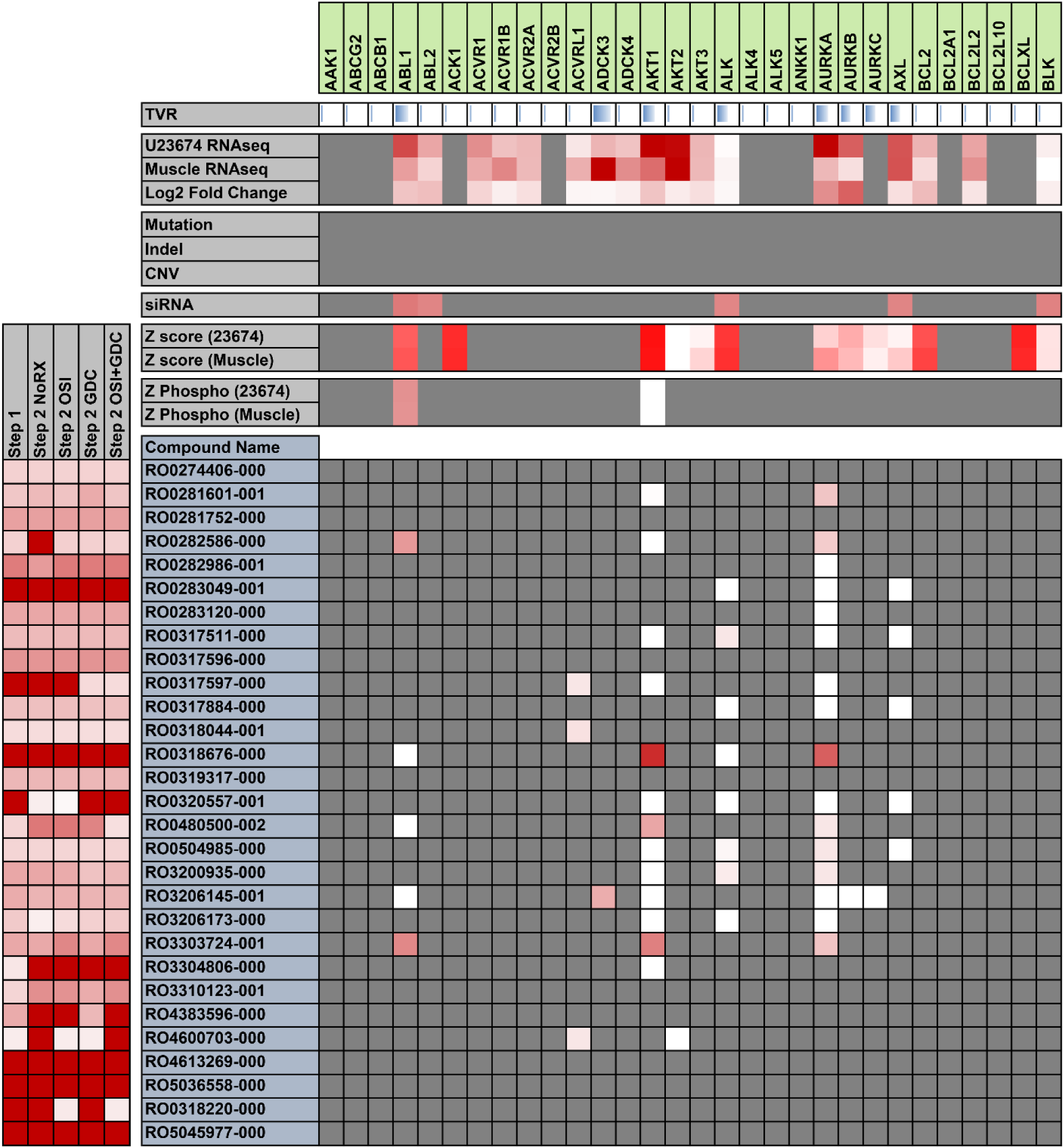
Heat map of joint Roche Orphan Kinome chemical screen, RNA-seq, siRNA, and phosphoproteomics results from the U23674 Probabilistic Target Inhibition Map (PTIM) rewiring experiment. Due to the large number of compounds and protein targets, only a limited scope of compounds and targets is shown here (For full data, see Supplemental Tables 14). Bright red indicates high sensitivity values, gradating down to white meaning low sensitivity. Gray indicates no interaction or no available data.

**Supplemental Figure 11.**
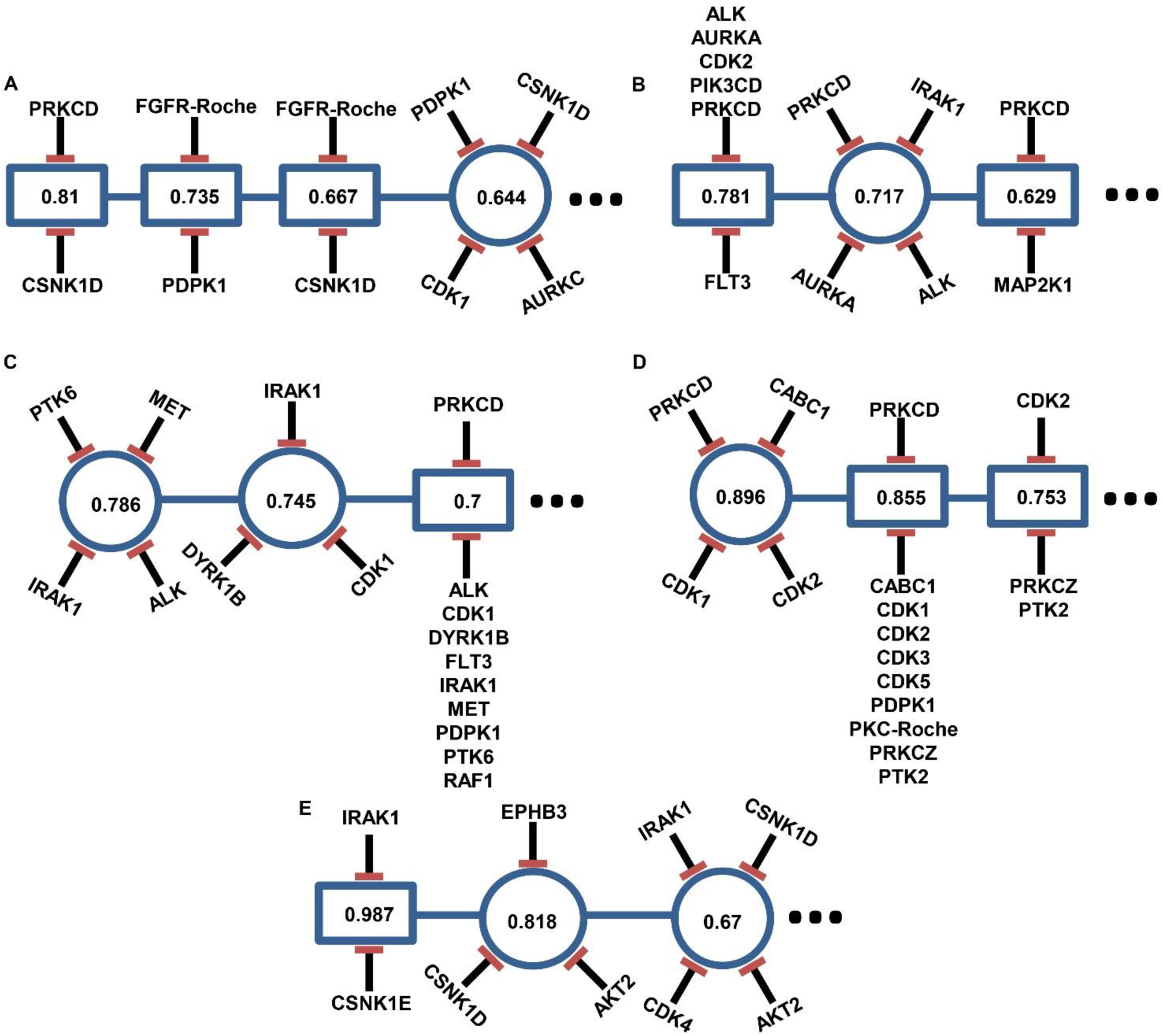
Probabilistic Target Inhibition Map (PTIM) models from U23674 experimental rewiring data. Values in the center of PTIM blocks represent expected scaled sensitivity following inhibition of associated block targets. (**A**) Untreated initial culture PTIM. (**B**) Untreated secondary culture PTIM. (**C**) OSI-906-treated rewire PTIM. (**D**) GDC-0941-treated rewire PTIM. (**E**) OSI-906 + GDC-0941-treated rewire PTIM.

**Supplemental Figure 12.**
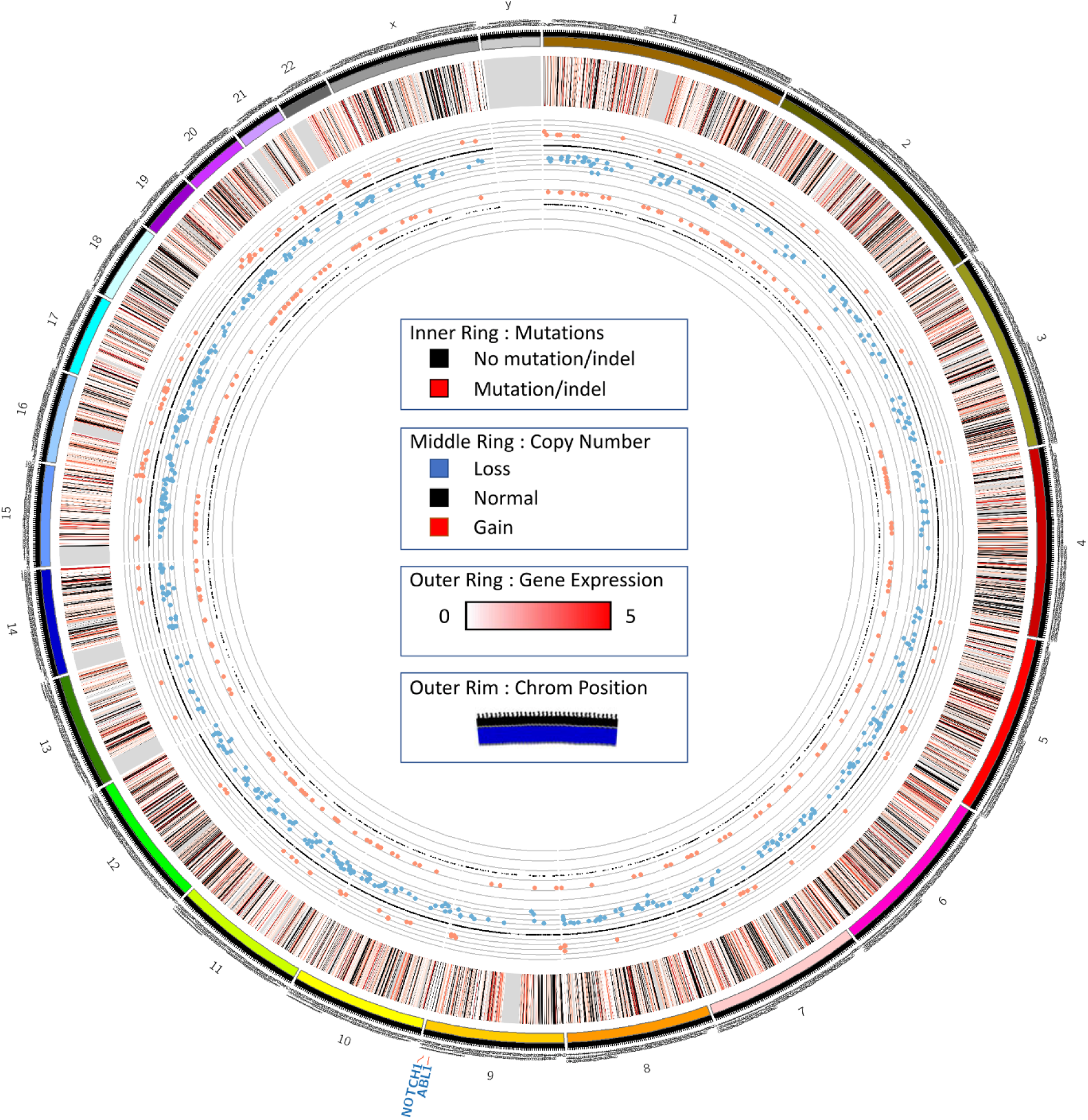
Circos plot of PCB490 RNA sequencing and exome sequencing data. *ABL1* and *NOTCH1* are identified as both mutated and amplified, though both variations were also identified in the matched germline sample. The outermost data circle represents log_2_-scaled gene expression [log_2_(expression+1), low expression (white) to high expression (red), with missing values colored black]. The middle circle represents genes with identified mutations or indels (red) or lack thereof (black). The innermost circle represents copy number variations (red is amplification, blue is deletion, black is no variation).

**Supplemental Figure 13.**
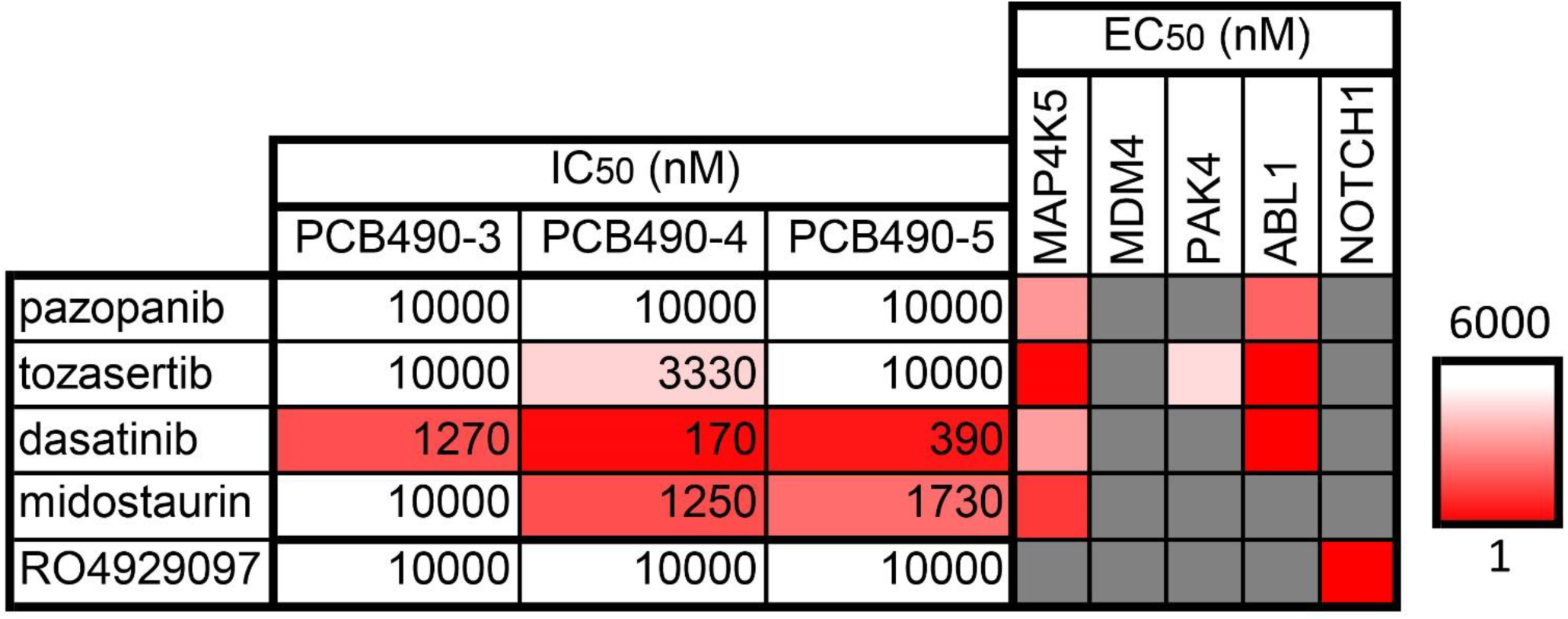
Heat map of IC_50_ and EC_50_ values for Pediatric Preclinical Testing Initiative Version 3 drug screen compounds inhibiting mutated and expressed targets in PCB490. Red in the IC_50_ and EC_50_ tables indicates low IC_50_ and EC_50_ values, respectively. No single target or combination of targets showed uniform efficacy across all PCB490 cultures, suggesting variations alone or in conjunction with transcriptome sequencing would have identified actionable therapeutic targets.

## SUPPLEMENTAL TABLES

**Supplemental Table S1.** Merged GSK Screen Data - U23674

**Supplemental Table S2.** GSK Screen Data IC50 Data - U23674

**Supplemental Table S3.** Roche Screen Merged Data - U23674

**Supplemental Table S4.** Roche screen hit references

**Supplemental Table S5.** Roche screen IC50 data - U23674

**Supplemental Table S6.** PPTI screen IC50 data - U23674

**Supplemental Table S7.** PPTI Screen Merged data - U23674

**Supplemental Table S8.** Exome Sequencing Data - U23674

**Supplemental Table S9.** RNA Sequencing Data - U23674

**Supplemental Table S10.** RAPID siRNA Screen data - U23674

**Supplemental Table S11.** Rapid Screen vs Drug screen - U23674

**Supplemental Table S12.** Phospho Screen data - U23674

**Supplemental Table S13.** Combination Index Values - U23674

**Supplemental Table S14.** Rewiring Screening Data - U23674

**Supplemental Table S15.** V3 Drug Screen data - PCB490

**Supplemental Table S16.** Roche Drug Screen data - PCB490

**Supplemental Table S17.** Exome Sequencing Data - PCB490

**Supplemental Table S18.** RNA Sequencing data - PCB490

**Supplemental Table S19.** Druggable Exome Targets - PCB490

**Supplemental Table S20.** EPS Model V3 Drug Screen Data

**Supplemental Table S21.** PPTI Drug Screening Data - UPS

